# Evolution under competition increases phytoplankton production by reducing the density-dependence of net energy fluxes and growth

**DOI:** 10.1101/2024.09.25.614915

**Authors:** Charlotte L. Briddon, Ricardo Estevens, Giulia Ghedini

## Abstract

Competition can drive rapid evolution but forecasting how species evolve in communities remains difficult. Life history theory predicts that evolution in crowded environments should maximise population production, with intra- and inter-specific competition producing similar outcomes if species compete for similar resources. Despite its appeal, this prediction has rarely been tested in communities. To test its generality and identify its physiological basis, we experimentally evolved four species of marine phytoplankton (spanning three orders of magnitude in cell size) alone or together in a community for 4.5 months. We then quantified changes in their metabolism, demography, and competitive ability at two timepoints (∼60 and 120 generations) in common garden experiments. One species was outcompeted during the evolution experiment. For the other three, we found the same evolutionary outcome: species evolved greater biovolume production regardless of competition treatment but did so either by increasing max. population size or individual cell size. Biovolume production increased because of the differential evolution of photosynthesis and respiration under intense competition. These metabolic changes meant that intraspecific competition decreased and cells maintained higher rates of net energy production and growth as populations neared the stationary phase. Overall, these results show that intra- and inter-specific competition influence physiological and population parameters similarly in species that compete for essential resources. Life history theory thus provides a valuable base for predicting how species evolve in communities, and our results show how these predictions connect with the evolution of metabolism and competitive ability.

## INTRODUCTION

Biodiversity change can drive rapid evolution because species introductions or losses alter competition regimes (1). However, which and how species traits evolve in response to competitive interactions remains difficult to anticipate (2, 3).

Macroevolutionary patterns indicate that traits should diverge when species compete for similar resources (4). The evolved trait differences should promote coexistence by increasing niche differences (5). But theory also suggests that, when species compete for essential resources, competitive ability may be more likely to evolve because opportunities for niche differentiation are limited (6, 7). Recent empirical work confirms these theoretical predictions: evolution in response to competition can result in a more efficient use of resources (e.g., evolved lower minimum resource requirement; (8, 9)), thus increasing competitive ability (6). However, requirements for different resources are linked through complex metabolic pathways (thus are not independent), and growth and fitness might change nonlinearly with minimum resource requirements (10, 11). So, assessing which specific resource traits evolve can indicate the driver of selection but does not necessarily facilitate predictions of evolution (12).

Density-dependent processes of population regulation are nearly ubiquitous in nature and link ecological and evolutionary dynamics (13). Thinking about evolution under competition in terms of changes in density-dependence can thus help generalise predictions across species and connect ecology with life history theory (14). An advantage of life history theory is to provide a clear prediction about the quantity that should be maximised under competition. Originally, MacArthur predicted that evolution in a stable and crowded environment should maximise the equilibrium size or carrying capacity (K) of a population (15, 16). Subsequent refinements of these predictions show that, when the environment fluctuates (i.e. there is variation in the strength of density-dependence), both population intrinsic growth rate (r) and K are maximised – thus competition should select for greater population production (17, 18). Despite its appeal, there are few empirical tests of this prediction in response to intraspecific competition (19, 20), and almost none under interspecific competition (21).

The lack of tests may partly be explained by the controversy regarding ideas of r-K selection (22). The assessment of the parameter K in particular poses some challenges because, in the r-K formulation of the logistic growth model, K is treated as an independent biological parameter but in fact K is not independent from r (22, 23). This issue can be resolved by using the original formulation of Verhulst’s growth model (24), with explicit intraspecific competition coefficient, because it emphasises the underlying demographic processes of growth and energy use (even though both models describe the same dynamics) (13, 23, 25).

We test the predictions of life-history theory under intra- and inter-specific competition and explain these evolutionary outcomes by connecting population parameters with metabolic traits (i.e., photosynthesis, respiration and net energy production as their difference over 24 hours). Demographic responses should covary with metabolism since energy fluxes influence growth and competitive ability (26–28). Since metabolic rates can evolve rapidly (29, 30), their assessment can help identify the physiological basis of evolved changes in competitive ability and demography.

We base our assessment on marine phytoplankton because these species have rapid generation times and high population densities with substantial capacity for rapid evolution (29), with adaptation generally occurring within a few weeks (31, 32). We evolved four marine phytoplankton species either alone (intraspecific) or together in a community (interspecific competition). We used species that span ∼ three orders of magnitude in cell volume because body size strongly influences metabolic rate and interspecific interactions (33–35). We then quantified changes in metabolism, demography and morphology of the evolved populations at two different time points in common garden experiments (after 9 and 17 weeks of evolution, corresponding approximately to 60 and 120 generations). Our goals are to 1) determine if evolution under intra- and inter-specific competition produces the same evolutionary outcomes, i.e. maximises population production as predicted by life history theory; 2) identify the physiological changes that underpin population-level responses, including the evolution of population parameters (growth rate, sensitivity to intraspecific competition); 3) test if species that differ in size (hence metabolic and competitive traits) show the same evolutionary trajectories and underlying mechanisms of evolution. Linking the demographic and physiological processes that alter population production under different forms of competition can help clarify current theoretical expectations for how species should evolve within ecosystems.

## MATERIALS AND METHODS

To test the effects of competition on the evolution of metabolism, morphology and demography, we used four species of marine phytoplankton (*Amphidinium carterae* RCC88*, Dunaliella tertiolecta* RCC6*, Tisochrysis lutea* RCC90 and *Nannochloropsis granulate* RCC438) acquired from the Roscoff Culture Collection, France. These species belong to different taxonomic groups and span almost three orders of magnitude in cell volume (*Amphidinium* = 949 ± 15 µm^3^; *Dunaliella* = 624 ± 4 µm^3^; *Tisochrysis* = 86 ± 0.66 µm^3^; *Nannochloropsis* = 19 ±0.17 µm^3^). To determine if evolution under intra- or inter-specific competition produces the same evolutionary outcomes we evolved these species either alone (“monoculture”) or together (“polyculture”) for 4.5 months (Phase 1: Evolution experiment ∼120 generations). In Phase 2, we used two common garden experiments (completed in weeks 9 and 17, corresponding to generation ∼60 and ∼120 respectively) to quantify trait evolution as explained in detail below.

### Phase 1: Experimental Evolution

We used dialysis bags cut to approximately 25 cm in length (MWCO 14 kDa, pore size 25 Angstrom, Dialysis membrane Membra-CEL, Carl Roth, Germany) to evolve each species under intra- or inter-specific competition. The bags allow competition for light and nutrients but maintain physical separation between species, thus we could phenotype each independently in the common garden experiments (similar to (21)). Before starting the experiment, we grew a large volume (∼2L) of each species (strain) in isolation and used this same well mixed culture to fill each bag of that species.

For the intraspecific treatment (mono), we set up three replicate beakers per species. Each beaker contained four dialysis bags, each filled with the same conspecific strain surrounded by enriched seawater medium (f/2 media prepared from 0.2 µm filtered and autoclaved natural seawater (36) containing no phytoplankton. This resulted in 12 dialysis bags split over three beakers for each species (Figure 1). For the interspecific treatment (poly), we set up 10 replicate beakers using the same design but each of the four dialysis bags was filled with a different species (Figure 1). Independently of the treatment, each dialysis bag had the same initial biovolume of 5.2 × 10^9^ µm^3^, filled to a volume of 45 ml with f/2 media. Each beaker was filled to 500 ml with fresh media to completely submerge the bags.

**Figure 1:**
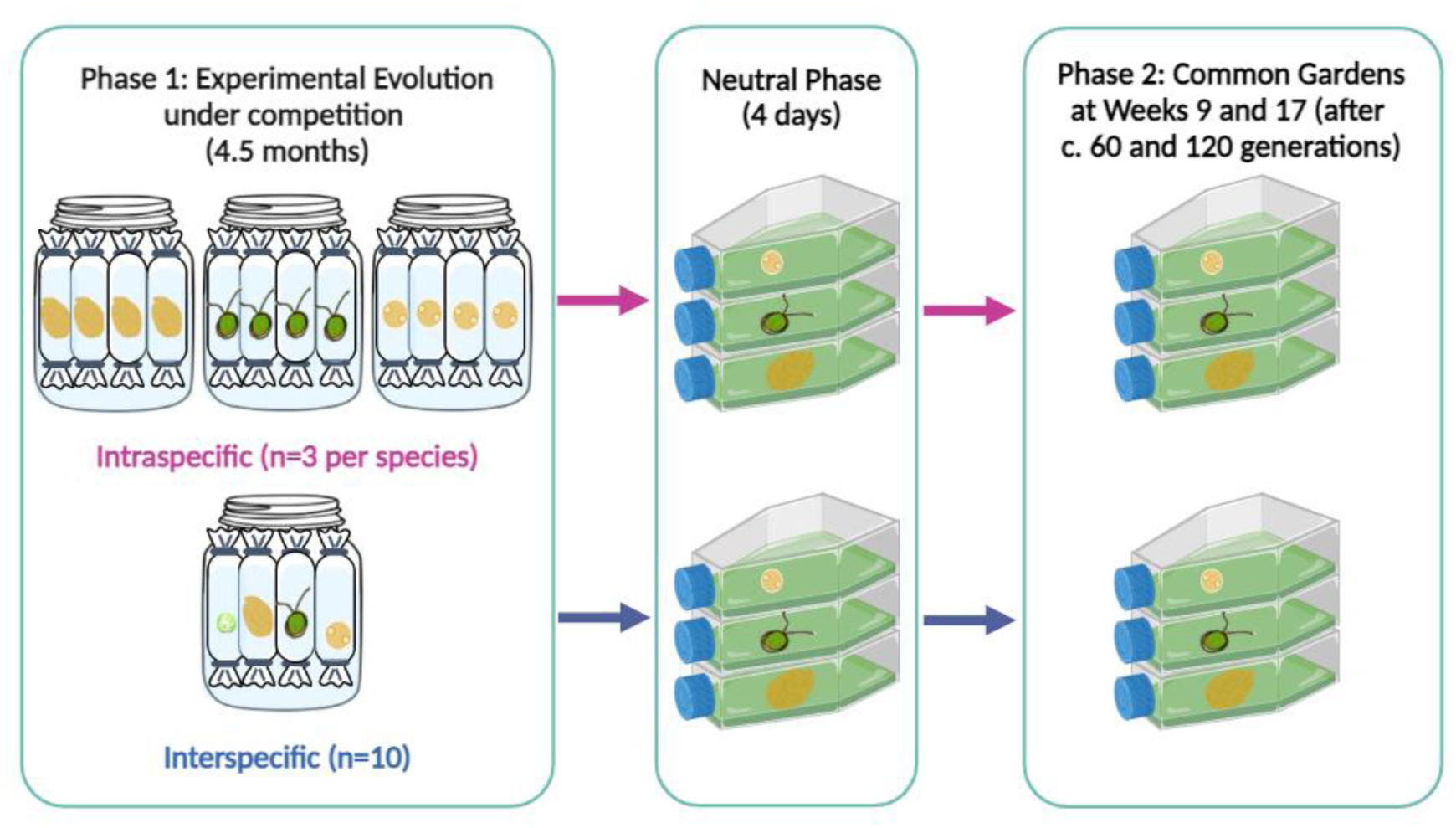
Schematic showing the experimental set-up of Phase 1 (Experimental Evolution) and Phase 2 (Common Gardens). During Phase 1, we evolved three species (*Amphidinium*, *Dunaliella*, *Tisochrysis*) either under intraspecific (monoculture) or interspecific competition (polyculture) for 4.5 months. At two time points (∼60 and 120 generations), we used common garden experiments (Phase 2) to assess the evolution of each species. Before each common garden, we had a phase of neutral selection (4 days) to remove any plastic response.

Once a week, we transferred a set volume of 25 ml from each dialysis bag to a new sterilised bag and beaker. The contents of the bag were topped up with 20 ml of fresh media. We also replaced the media in the beaker. All beakers were bubbled daily with filtered atmospheric air (0.22 µm; Minisart, Sartorius, Göttingen, Germany) for 15 min every hour. For weeks 5-10, we reinoculated all bags with a lower set volume of 15 ml (instead of 25 ml) due to the high cell density to avoid nutrient limitation. After week 7, the populations of *Nannochloropsis* began to decline, particularly in the interspecific (poly) treatment, so this species was not included in the second common garden (week17) and therefore was removed from all analyses. Light intensity was set at 60 µmol m^-2^s^-1^ with a 12:12 light:dark schedule using low-heat 50W led flood lights (Surface Luminária LED 230V, Robert Mauser, Portugal). The temperature was maintained at 20 ± 1℃.

### Phase 2: Common Garden (CG) experiments

After 9 and 17 weeks of evolution (∼60 and 120 generations respectively), we phenotyped the evolved lineages from the mono and poly treatments for morphological, metabolic and demographic changes to determine how these traits evolved. We tested two bags from each beaker for the mono treatment (n = 3 per species as the two bags were not independent) and all bags for the poly treatment (n = 10 for each species). Before beginning each CG experiment, we placed a 10 ml sample of each bag in a neutral environment (i.e. cell culture flask filled up to 100 ml with f/2 media) for four days (∼ 4 generations) to remove any environmental conditioning. For both CGs, we then inoculated an equal biovolume of 2.5 × 10^9^ µm^3^ into 250 ml culture flasks, filled up to 100 ml of media. Both CGs lasted for 25 days, until the samples had reached carrying capacity. For each sampling day, we collected 10 ml from each culture flask for analysis (see details below) and replaced it with 10 ml of fresh media. For both CGs, we measured on day 1, 2, 4, 5, 7, 9, 11, 14, 16, 18 (week17 only), 21 and 25.

### Cell size, densities and biovolume

On each sampling day of the CG, we measured cell size (µm^3^), cell density (cells/µl) and biovolume (µm^3^/µl; calculated by multiplying cell density and average cell size for each replicate). Size and density were obtained from photos taken with light microscopy at 400x magnification (Olympus IX73 inverted microscope) after staining a 1 ml sample of each sample with 1% Lugol’s iodine. Ten µl of the sample were loaded onto a Neubauer counting chamber (Marienfield, Germany) and 20 evenly spaced photographs were taken around the counting frame. All images were analysed using ImageJ and Fiji software (version 2.0) (37) to quantify the different morphological characteristics and cell densities. The software recorded length and width of each cell, and we then estimated cell volume by assigning either a prolate spheroid shape (*Amphidinium*, *Dunaliella*) or spherical shape (*Tisochrysis*) (38).

### Metabolic Rates

In conjunction with biovolume and density data, we measured photosynthesis and respiration rates to identify the physiological changes that underpin population-level responses. For each lineage, we tracked changes in oxygen saturation in 5 ml vials using established protocols (21). Prior to all experiments, the respirometry system (PreSens Sensor Dish Reader, SDR; PreSens Precision Sensing, Germany) was calibrated using 0% and 100% oxygenated seawater. To prevent carbon limitation, we added 50 µl of sodium bicarbonate stock (a final concentration of 2mM sodium bicarbonate) to each sample vial before measurements began. The samples were measured for 20 minutes in the light to quantify photosynthesis, followed by 40 minutes of darkness to determine respiration rates. As the cultures were not axenic, sixteen blanks were filled with spent media with no phytoplankton cells to account for the background bacterial activity (spent media was obtained by centrifuging samples at 5,000 rpm for 10 minutes to separate the algal cells from the supernatant). The photosynthesis and respiration rates (VO_2_; μmol O_2_/min) of each sample were calculated as:

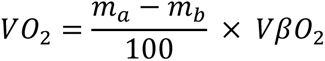

where m_a_ is the rate of O_2_ saturation change of the sample (min^-1^), m_b_ is the mean O_2_ saturation across all blank samples (min^-1^), V is the sample volume (0.005 L) and βO_2_ is the O_2_ capacity of air-saturated seawater at 20℃ and 35 ppt salinity (225 µmol O_2_/L) (39). The first three minutes of measurements under the light were discarded to allow acclimation; similarly, respiration rates were calculated after 20 minutes of dark when the O_2_ levels demonstrated a linear decline. Subsequently, we converted the photosynthetic and respiration rates (µmol O_2_/min) to calorific energy (J/min), using a conversion factor of 0.512 J/µmol O_2_ to estimate the energy production and consumption respectively (40). These calculations were completed using the LoLinR package (41) in R (version 4.3.2) (42).

### Data Analysis

The statistical analyses were completed on the data collected during the CGs to assess evolutionary differences instead of plastic responses. All data was analysed using R (version 4.3.2) (42) and Rstudio (43) using the packages nlme (44), lme4 (45), emmeans (46), car (47), plyr (48) for analyses and ggplot2 (49) (50) for creating plots. For all statistical analyses, any interactions between the covariates were removed when p > 0.25. All figures with means show the least square means with a 95% confidence interval using a Tukey p-value adjustment unless stated otherwise.

**a. Changes in r and K.** We calculated the maximum rate of increase (r_max_) and the maximum predicted value (i.e. carrying capacity, K) of biovolume (μm^3^/μl) or cell density (cells/μl) for each replicate to determine how demography evolved under each competition treatment (mono, poly) and common garden experiment (week 9, week 17). We implemented the same approach used in (51, 52). Firstly, we fitted four growth models to each replicate using non-transformed growth data (biovolume or cell density).The models were: a logistic-type sinusoidal growth model with lower asymptote forced to 0 (e.g. three-parameter logistic curve), a logistic-type sinusoidal growth model with non-zero lower asymptote (e.g. four-parameter logistic curve), a Gompertz-type sinusoidal growth model (e.g. three-parameter Gompertz curve) and a modified Gompertz-type sinusoidal growth model including population decline after reaching a maximum (e.g. four-parameter Gompertz-like curve including mortality). Secondly, we utilised the AIC (Akaike information criterion) to determine the best fitting models for each lineage with successful convergence, which we used to estimate the maximum predicted value (K) of biovolume or cell density for each replicate. From the first derivative we extracted the maximum rate of increase (r_max_). Finally, we used a linear model to test for differences in K and r_max_ between species and competition treatments. To avoid pseudoreplication for the mono treatment, the data from the two samples (bags) collected from the same beaker were averaged, resulting in n = 3 for the subsequent analyses (unless otherwise stated). In the main text, we report the test on the percentage change (calculated from the difference in K or r_max_ at week 17 relative to week 9) using a linear model that includes species identity (*Amphidinium, Dunaliella, Tisochrysis*) and competition treatment (mono, poly) as factors. We report the figures and analyses on differences in absolute values of r_max_ and K in the supplement (Figure S2-S3), where we test for differences in population parameters of each species separately using a linear model that has initial biovolume (or cell density) as covariate and competition treatment (mono, poly) and experiment (week 9, week 17) as factors, including their interactions.

From the best-fitting growth model for each replicate we also extracted the population growth rate (cells/μl) on each day of the common gardens. We then analysed these growth rates from day 8 onwards (i.e. the declining phase) to identify evolved changes in growth rates as species approach stationary phase. We used a linear mixed effect model that includes growth as a response variable (log_10_-transformed), CG Day, experiment, competition treatment and species (including their interactions), and sample code as a random effect.

To visualize differences in growth trajectories (Figure S1), we fitted the same models described above to all replicates within a treatment (species, competition treatment and common garden). We then plot the best-fitting model based on AIC for each treatment combination.

**b. Changes in competitive ability.** To better understand the demographic changes in r_max_ and K, we fitted biovolume data with a logistic growth model with explicit r-α formulation (24, 25). With this model we can determine how the strength of intraspecific competition α_ii_ evolved over time as a function of competition type. We fit the model to biovolume data of each replicate (B, μm^3^/μl) because biovolume is the quantity that is more comparable and evolves in the same way among species:

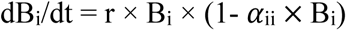

Subsequently, we used a linear model to determine how α_ii_ changes as a function of intrinsic rate of increase (r), experiment (week 9, week 17), competition treatment and species. Interactions were removed when p > 0.25 so the final model took the form of: *α*_ii_ ∼ r + experiment + treatment + species + r × species + experiment × treatment + experiment × species + treatment × species. For this analysis we used all replicates (not averaging between beakers) because we were interested in the evolution of density-dependence and its relationship with r within each population. We excluded one *Amphidinium* sample from the polyculture treatment that had an *α*_ii_ three times that of all other *Amphidinium* samples (3.6e-06 vs ∼1.2e-06), due to unusually slower growth in the first days of the common garden.

**c. Metabolic rates.** To explore the physiological basis of the demographic responses, we tested for changes in the density-dependence of net energy. First, we estimated *per capita* rates of photosynthesis or respiration by dividing the population rate of each replicate by its total population size (cells in 5 ml). We then estimated *per capita* net energy production over a 24-hour period (J/day/cell) as 12 hours of energy produced through net photosynthesis minus 12 hours of respiration. We removed day 0 (as no oxygen rates were measured on that day), as well as day 1 and day 2 because many respiration rates were negative (the biovolume was low so respiration rates were often not different or shallower than blanks). We had to remove an additional 18 data points that had negative values (16 for respiration rates and 2 for net energy). This left us with 713 datapoints in total. On this dataset, we fitted a linear mixed-effect model with *per capita* net energy production (log_10_-transformed) as a response variable, population biovolume (log_10_-transformed) as a covariate, experiment, competition treatment and species as factors, and sample code as a random effect.

To understand what was driving the change in net energy, we compare *per capita* photosynthesis and respiration rates during the exponential (day 4) and stationary phase (day 25). We use a simple linear model with *per capita* rates as the response variable (log_10_-transformed); species identity (*Amphidinium, Dunaliella, Tisochrysis*), “condition” and growth phase (exponential vs stationary) as predictors including their interactions. Note that here “condition” combines both Experiment and competition treatment (i.e., week9_mono, week9_poly, week17_mono, week17_poly) to avoid overfitting the model (and since we have already seen from the analysis above that competition had small effects).

**d. Changes in cell size.** To determine how cell size evolved under competition, we first compared cell sizes on day 0 of each common garden with the ancestral cell size of each species. We used a linear model with the cell size (log_10_-transformed to meet normality assumptions) as the response variable and species (*Amphidinium, Dunaliella, Tisochrysis*) and condition (ancestor, week9_mono, week9_poly, week17_mono, week17_poly) as factors, including their interactions. Then we explored how size changed during common garden experiments for each species separately since they show different trajectories. We used a linear model that included competition treatment, experiment and common garden day as factors, and their interactions. The figure shows the measured cell sizes (the average cell size of each replicate) and the estimated marginal means with lower and upper confidence intervals; we also include the mean cell size of the ancestors (measured on the first day of the evolution phase) for comparison.

## RESULTS

### Population biovolume production increases in response to competition

All three species evolved greater population production at week 17 (relative to week 9) by increasing biovolume carrying capacity (K) at no expense of max. growth rates (r_max_) (Figure 2a-b; Figure S1-S2). Each species increased its max. biovolume by approximately 20% and the strength of this response was similar between intra- and inter-specific competition (Table S1, Table S2). For one species (*Tisochrysis*), the populations evolved in monoculture had higher biovolumes than those evolved in polyculture at both timepoints (Figure S2). *Tisochrysis* was also the only species to show a change in the max. rate of biovolume production (r_max_) which increased from week 9 to week 17, and more so in the polyculture treatment (Figure 2a, Figure S2, Table S2).

**Figure 2:**
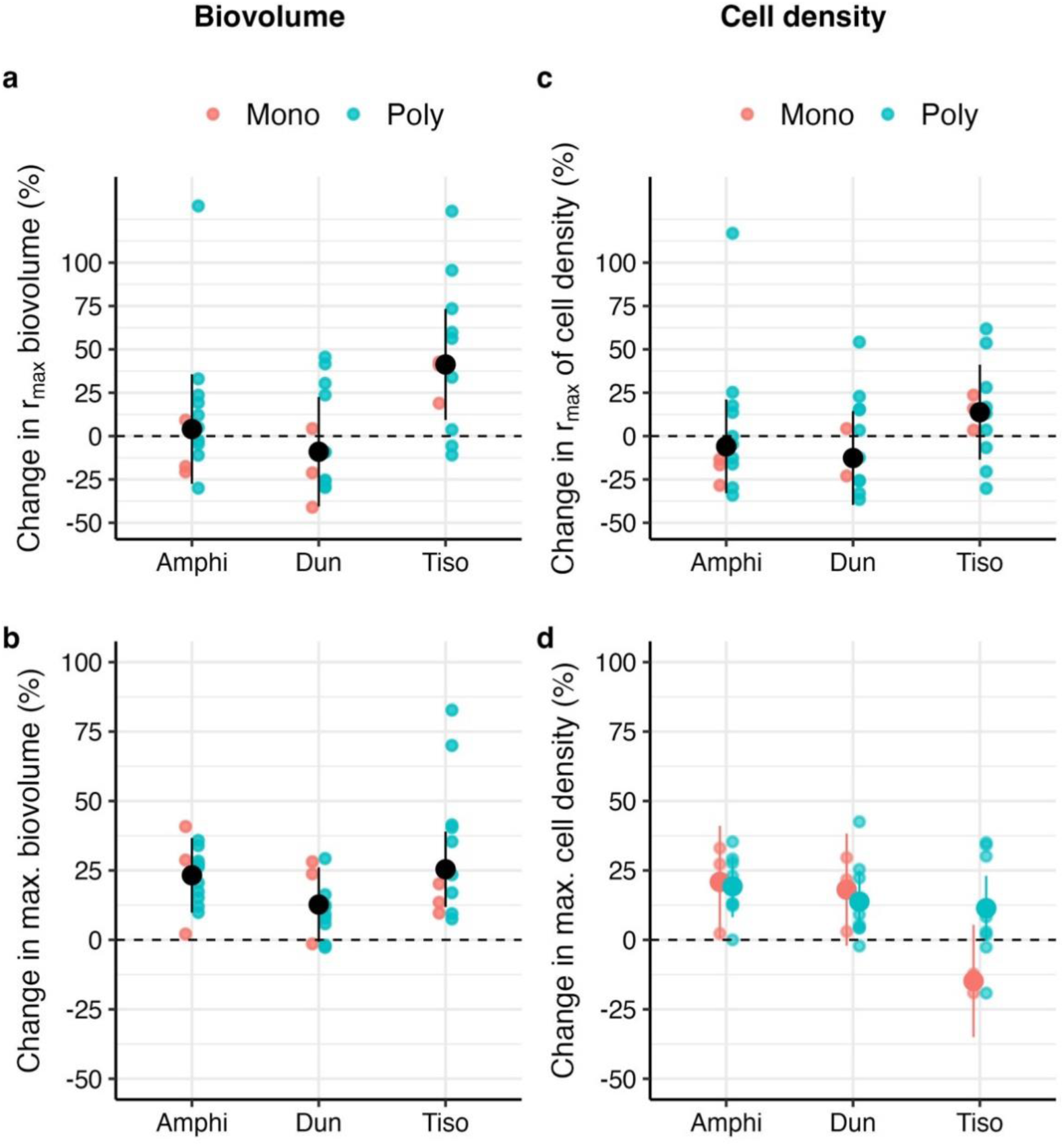
Percentage change in the maximum rates of increase (r_max_) and maximum values (K) of biovolume (a, b) and cell density (c, d) from the first to the second common garden (positive values indicate an increase, negative values a decrease from week 9 to week 17). All species increased the maximum biovolume (b) and two out of three also increased their cell density (d). *Tisochrysis* showed different changes in cell density depending on the competition treatment. Maximum growth rates did not change, either for biovolume (a) or cell density (c), except for *Tisochrysis* that evolves faster max. rates of biovolume production. Refer to Table S1 for the model outputs.

Biovolume carrying capacity increased because populations increased their max. population size in two species (*Amphidinium*, *Dunaliella*), and because cell size increased in the third species (*Tisochrysis*; Figure 2c-d; Figure S3-S5). The observed increases in cell densities were similar regardless of the competition history (Figure 2d), i.e. whether the species evolved with intra- or inter-specific competitors, with one exception: the max. cell density of *Tisochrysis* declined over time in monoculture and slightly increased in polyculture (albeit this increase was not significant) (Figure S3, Table S3). Max. population growth rates did not change for any of the species or treatments (Figure 2c, Figure S3).

In general, all three species reduced their cell sizes in comparison to the ancestors, at least when considering the size at the start of common garden experiments (Figure S4; Table S4). But cell size tended to increase as the CG progressed with no clear differences between competition treatments (Figure S5, Table S5). In the second common garden, *Tisochrysis* showed a marked increase in cell size from day 4 onwards (Figure S5) which explained why this species reached greater biovolume without increasing cell densities.

### The evolution of reduced density-dependence

To better understand how species increased their biovolume production, we assessed changes in density-dependence. All species reduced their sensitivity to intraspecific competition over time (experiment × species: F_2, 81_ = 4.66, p = 0.012; Figure 3a; Table S6); in other words, species improved their intraspecific competitive ability as *α*_ii_ declined from week 9 to week 17, thus reducing the density-dependence of biovolume growth (with significant differences for *Amphidinium* and *Tisochrysis* (p < 0.0001), but marginal for *Dunaliella* (p = 0.0948)). At both timepoints, we also find that intraspecific density-dependence was stronger in the populations evolved in polyculture compared to those evolved in monoculture (competition treatment effect: F_1,81_ = 8.89, p = 0.004; Table S6).

**Figure 3.**
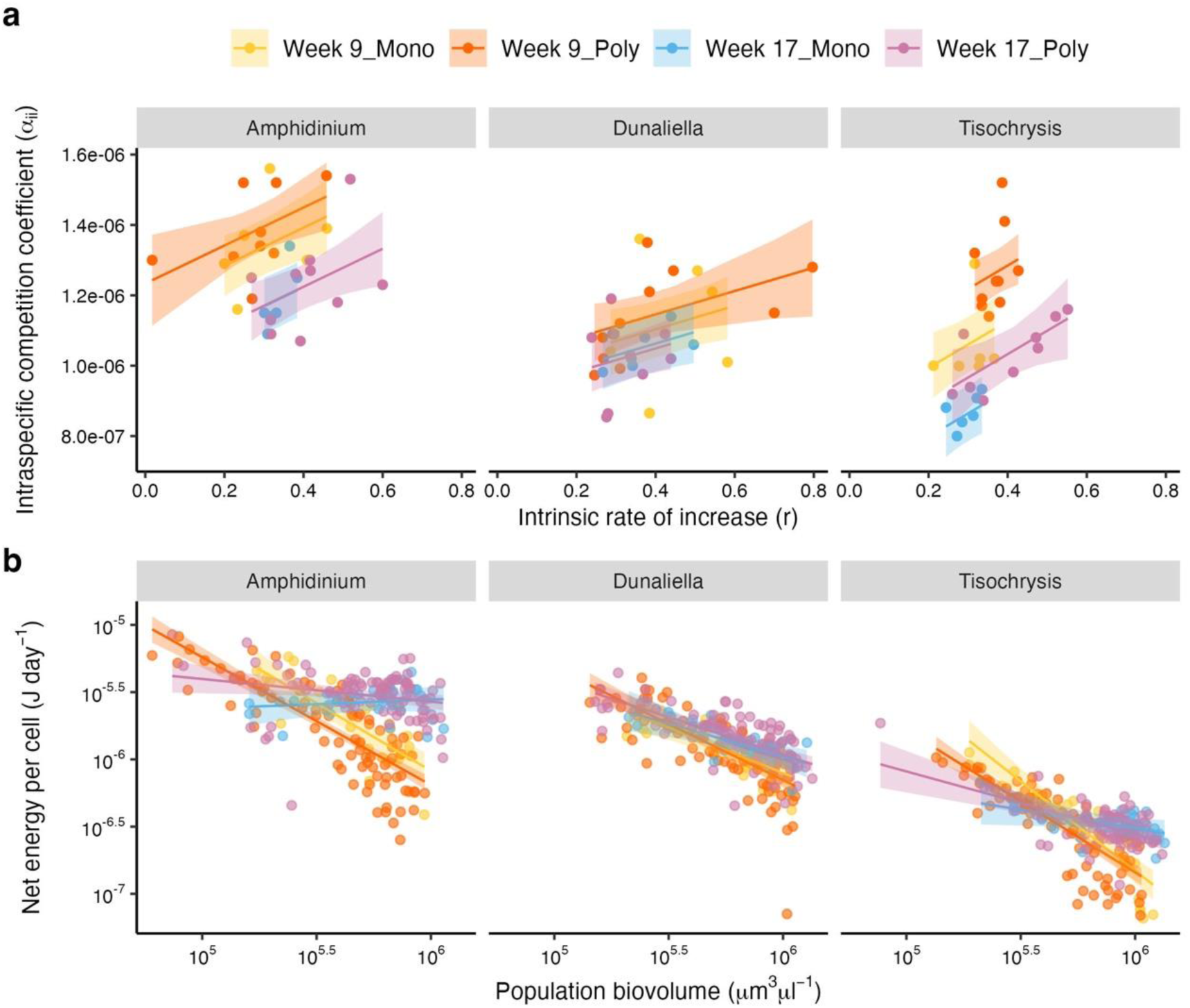
(a) The strength of intraspecific density-dependence (*α*_ii_ calculated on biovolume data) declines over time (from week 9 – warm colours – to week 17 – cold colours) across all species, although this decline is weaker for *Dunaliella* (posthoc comparison between week 9 and week 17: p = 0.09 for *Dunaliella*). Independently of time, density-dependence was stronger in populations from polycultures than monocultures. (b) Simultaneously, all species evolved a weaker density dependence of *per capita* net energy production as slopes were shallower in week 17 compared to week 9 for each species. Across all species, populations evolved in monocultures reduced their density-dependence more (slopes were significantly shallower) than populations evolved in polyculture at week 17. Refer to Tables S6 and S7 for the slopes and model outputs.

Competitive ability coevolved with changes in energy fluxes. All populations evolved a weaker density-dependence of net energy production at week 17, so that cells sustained higher net energy fluxes as their populations approached stationary phase (log_10_(biovolume) × experiment × species interaction: F_2,645_ = 11.1, p = <0.001) (slopes were shallower in week 17 compared to week 9 for each species; Figure 3b; Table S7). Across all species, we also found that populations evolved in polyculture tended to reduce their density-dependence less than those evolved in monoculture (had steeper a density-dependence of net energy in week 17, log_10_(biovolume) × experiment × treatment interaction: F_1,645_ = 2.9, p = 0.08), in agreement with the higher intraspecific competition coefficients observed above.

Increases in net energy were primarily driven by higher *per capita* photosynthetic rates, while we did not detect any change in respiration rates when populations were in stationary phase (Figure 4, Table S8). Photosynthesis rates were lower in the stationary phase than in exponential phase as expected from stronger resource limitation (Figure 4). But in stationary phase we observed a consistent pattern across species: photosynthesis rates increased from week 9 to week 17, although this response was significant only under interspecific competition for *Amphidinium* and *Dunaliella* (Figure 4, Table S8). There were no significant differences in photosynthesis rates in the exponential phase (except for *Dunaliella* that significantly increased photosynthesis from week 9 to week 17 in polyculture). Respiration rates were more variable during exponential phase, but all species evolved in polyculture reduced *per capita* respiration from week 9 to week 17 (Figure 4, Table S8).

**Figure 4.**
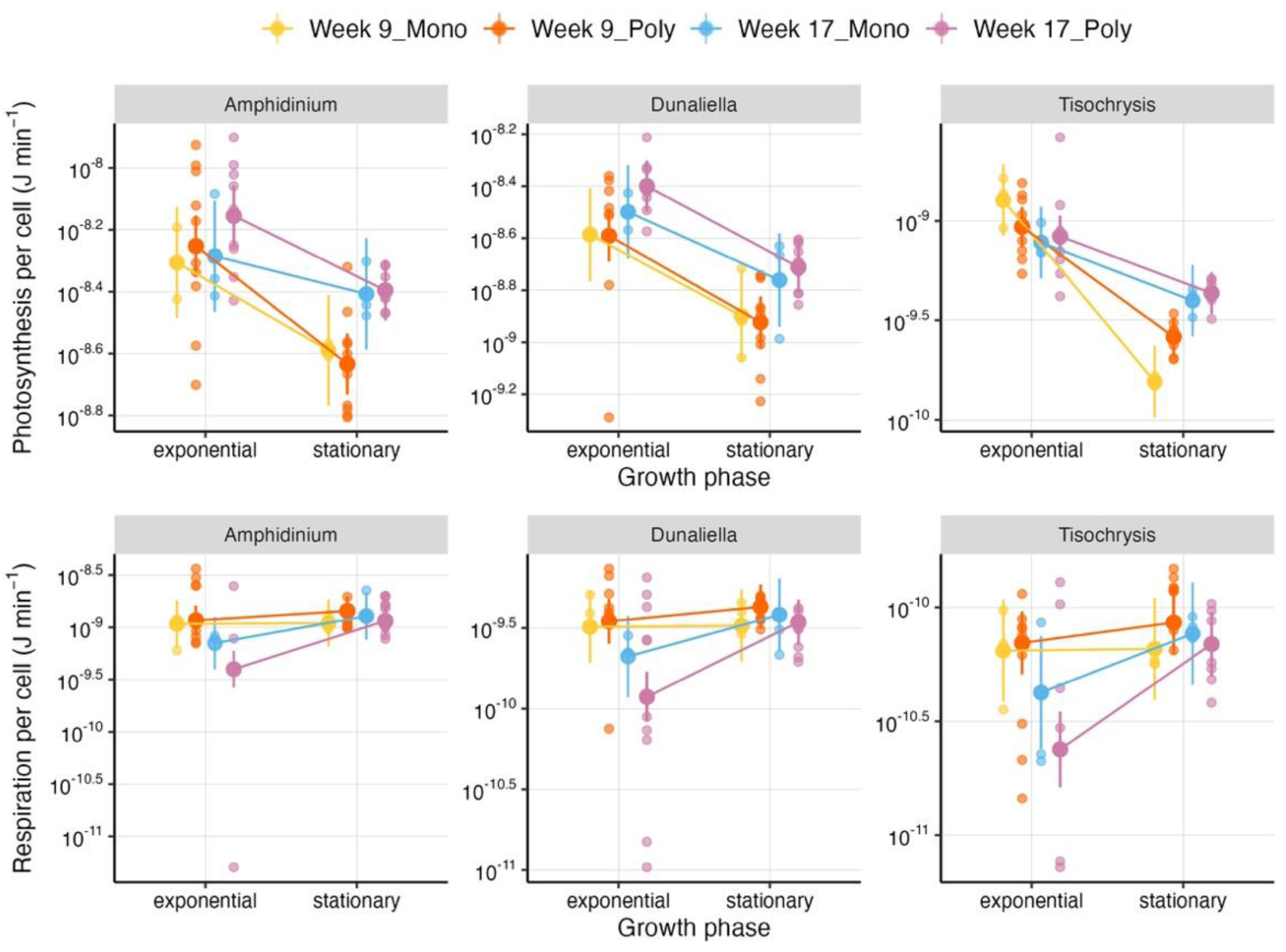
Reaction norms showing changes in *per capita* photosynthesis and respiration from exponential (day 4) to stationary phase (day 25) for the two common gardens and competition treatments. All species show an increase in photosynthesis rates from week 9 (warm colours) to week 17 (cold colours) in stationary phase. The strength of this increase varies between species and treatments (for *Amphidinium* and *Dunaliella* it is significant only in the polyculture treatment). Changes in exponential phase are not significant, except for *Dunaliella* between week 9 and week 17 in the polyculture treatment. Conversely, respiration rates do not differ in the stationary phase for any of the species but are significantly lower in the polyculture treatment at week 17 compared to week 9 (either in the mono or poly treatment). Refer to Table S8 for the model outputs.

### Increased net energy fluxes sustain faster growth in competitive environments

The reduced density-dependence of net energy, driven by the differential evolution of photosynthesis and respiration, meant that evolved populations grew faster as populations approached stationary phase (Figure S6). Population (and *per capita*) growth declined less steeply in week 17 compared to week 9 within each competition treatment. *Tisochrysis* evolved in monoculture was an exception, showing the opposite pattern (these populations achieved greater biovolume through changes in cell size and not changes in cell densities) (Table S9). At both time points (week 9 and week 17), growth declined faster for populations evolved in polyculture than in monoculture (Figure S6, Table S9), consistently with the more negative effects of interspecific competition on competitive ability (*α*_ii_) and net energy.

## DISCUSSION

Life history theory predicts that competition should increase population production (17, 18) but there are few empirical tests under interspecific competition. Therefore, the generality of this prediction remains unclear. MacArthur and Wilson (16) hypothesized that, in a community of species that compete for similar resources, interspecific competitors should compound the effects of intraspecific competition leading to similar evolutionary outcomes – which should be particularly true when species compete for essential resources. Our results validate these predictions as we found that all three species tested evolved greater population production in response to intra- and inter-specific competition, increasing biovolume carrying capacity (K) at no expense of max. growth rates (r_max_). Our species achieved this same outcome in two different ways, either by evolving greater maximum cell densities or larger cell sizes. This result suggests that evolution under competition increases population production in the form of biovolume and that different species might employ different strategies in maximising this quantity. We can speculate that smaller species (e.g., *Tisochrysis*, the smallest species we tested) might be more likely to increase biovolume through the evolution of larger cell sizes (rather than densities), but this can only be confirmed with further studies incorporating additional species.

Not only competition led to the same consistent outcome across species and competition treatments (i.e. greater total biovolume). We also found that the underlying physiological mechanisms were the same. Evolution reduced density-dependence so that cells sustained higher net energy fluxes and growth rates in crowded environments. Net energy results from the balance of photosynthesis and respiration over 24 hours. Therefore, net energy can increase either by reducing respiration costs or increasing energy production through photosynthesis. We found that the main driver of increases in net energy were photosynthetic rates which were upregulated across all species in the stationary phase after evolution. While we did not further explore how photosynthesis increased, cells can enhance productivity through chlorophyll synthesis and packing (51). Processes of energy intake/production and expenditure can thus respond differently to competition, not only in terms of plastic changes (53) but also evolutionary responses (54). This differential regulation of energy fluxes might be very important for competition and growth because organisms can increase energy gains without also increasing metabolic costs.

Indeed, while we cannot establish if physiology drives demography or the other way around, net energy and competitive ability (*α*_ii_) improved in parallel. Reduced sensitivity to competition evolved as a strategy to minimise competition from both intraspecific and interspecific competitors, as also found in plants (14), albeit we find that interspecific competition weakened this evolution. Improvements in competitive ability did not come at the cost of maximum growth rates; on the contrary max. growth rates improved in one species. Trade-offs between r_max_ and K are not always observed (23); if anything, r_max_ and K should be positively correlated (23, 55, 56). While we find weak to no evidence for changes in max. growth rates, growth improves in competitive environments: reduced density-dependence allows populations to maintain higher growth rates as population biovolume increases, thus accumulating more biovolume. Similarly, fly populations evolved in crowded conditions have a higher rate of *per capita* growth at high densities, while these improvements in performance do not manifest at low densities (57). Overall, it seems that competition maximises population production through changes in energy use that increase energy gains and the conversion of this energy into biomass (16, 54).

Why do max. growth rates not evolve? The adaptation of organisms to ‘harsh’ environments (such as intense competition) should result in a trade-up of r_max_ and K if traits are far from optimised, as should be in our experiment since none of the species experienced this experimental set up before (14, 17, 18, 20, 23, 58). Similarly, variations in selective pressures due to environmental fluctuations (e.g. transfers from low to high densities) should optimise both max. growth rates (i.e. growth when resources are abundant) and competitive ability (i.e. efficiency when resources are scarce) (16, 17, 59). Instead, only one species (*Tisochrysis*) improved r_max_, which was positively correlated with max. biovolume. Since the strength of selection on max. growth is linked to environmental conditions (56), stronger fluctuations between low- and high-density environments (e.g., more frequent or larger dilutions) could increase selection and result in the evolution of r_max_ across more species (6).

Evolutionary responses were very similar under both competition treatments. This similarity could be due to an experimental limitation: dialysis bags prevent species from being in direct contact, which can increase intraspecific competition at the expense of interspecific competition in the polyculture treatment. This dialysis bag approach was nonetheless essential to phenotype each species independently in common gardens (cell sorting was not a viable approach because population recovery and growth was extremely slow). Nonetheless, when species compete for essential resources, the identity of the competitor might be of little importance (16, 60). The metabolism and growth of phytoplankton seems primarily affected by the total amount of competition (determined by total biovolume) and less so by the specific composition of biovolume (i.e. the abundance of intra- and inter-specific competitors) (61).

Despite the overall similarity in evolutionary responses, we observed differences in the strength of responses driven by intra- and inter-specific competition. Reductions in density-dependence were weaker when evolution occurred in polyculture: these populations showed higher intraspecific competition coefficients (Figure 3a), stronger density-dependence of net energy (Figure 3b), and reduced growth compared to those evolved in monoculture (Figure S6). These effects were generally small and did not affect the main evolutionary outcome (increased biovolume production). Interspecific competitors might have weakened evolution in two opposing ways. The presence of multiple competitors might allow resource partitioning (62), thus reducing the selective pressure of competition on density-dependence relative to monoculture treatments (if total biovolume is the same) – which could explain why these populations evolved “less” in terms of density-dependence. Alternatively, if resources are more fully utilized in communities, interspecific competition might intensify resource depletion and reduce opportunities for niche diversification (which can alleviate density-dependence) (63, 64). The reduced fitness of species evolved in polyculture (i.e. faster decline in growth rates as populations approach stationary phase, Figure S6) would point to this latter explanation.

Phytoplankton cell size affects metabolic rates, resource uptake and competitive ability (65). Therefore, species of different sizes might evolve in different ways. Intra- and interspecific competition both increased population biovolume regardless of species size. Yet, the three species maximised biovolume through different life history strategies (by increasing max. population size or individual size), although we cannot establish if these strategies are driven by differences in size. We initially included a fourth species, *Nannochloropsis* (the smallest species), which went extinct halfway through the evolution experiment and first in the polyculture treatment. Smaller algae have a higher surface-to-volume ratio allowing for greater rates of resource uptake per unit of biomass and suffer less from self-shading (12); therefore, they should be strong competitors. However, small cells have lower storage capacity and poorer recovery from nutrient depletion (51, 66). The advantage of larger cells in terms of nutrient storage could explain why *Nannochloropsis* was competitively excluded and why *Tisochrysis* increased cell size in response to competition (33). The other two species reduced size in comparison to the ancestor (similarly to (21), but these species still had much larger volumes (61). Future experiments using species of varying sizes or with different experimental set-ups could provide further insights into whether cell size influences evolutionary responses to competition.

## CONCLUSIONS

In summary, our results validate the prediction of life history theory that evolution under density-dependence maximises population production (16, 60). We further show the common physiological mechanism that underpins these demographic responses and its connection with competitive ability. The differential evolution of photosynthesis and respiration weakened the density-dependence of net energy fluxes – in other words cells produced more net energy (and thus biomass) in crowded conditions. These physiological changes also meant that intraspecific competition declined (*α*_ii_) and growth rate increased. Our work shows that intra- and inter-specific competition lead to the same evolutionary outcomes in phytoplankton, suggesting that similar patterns might hold when species compete for essential resources. The concomitant assessment of the physiological and demographic traits that underpin density-dependence should facilitate the identification of the mechanisms that drive evolution. Determining these eco-evolutionary dynamics seems particularly important in face of rapid biodiversity and environmental change (67).

## ACKNOWLEDGMENTS

We would like to thank the attendees of the Gordon Research Conference Unifying Ecology Across scales 2024 for fruitful discussions about this work. We also thank the IGC’s Advanced Imaging Facility, especially José Marques for help with automated image analyses. GG was supported by a fellowship (LCF/BQ/PI21/11830001) from ‘‘la Caixa’’ Foundation (ID 100010434) and the European Union’s Horizon 2020 research and innovation programme under the Marie Sk1odowska Curie grant agreement no. 847648.

## AUTHOR CONTRIBUTIONS

CB performed the research, analysed data and wrote the paper. RE performed the research. GG designed the experiment, analysed data and wrote the paper.

