## Supplementary Information for "Evolution under competition increases phytoplankton production by reducing the density-dependence of net energy fluxes and growth"

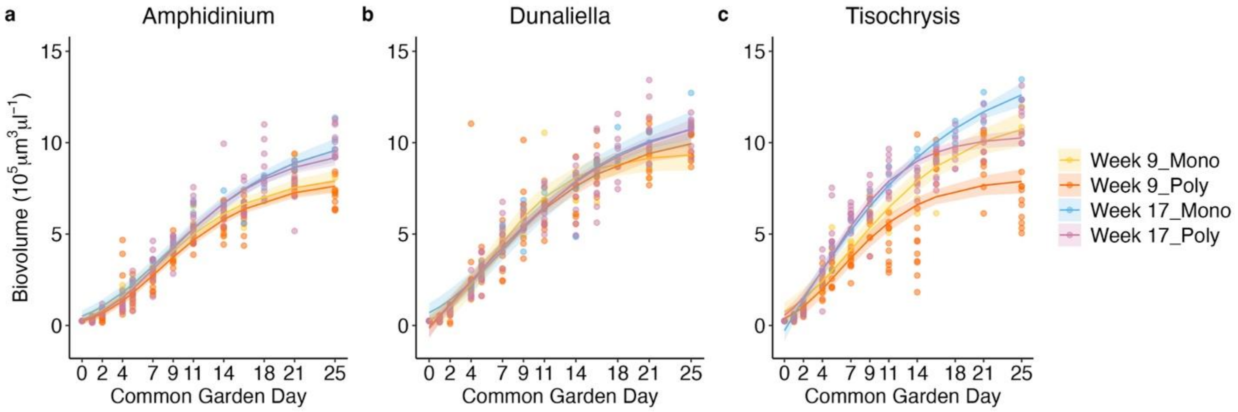

**Figure S1.** Biovolume carrying capacity increases over time across all species. These increases are similar between competition treatments, except for *Tisochrysis* populations that have higher biovolumes when evolved in monoculture compared to polyculture at both timepoints.

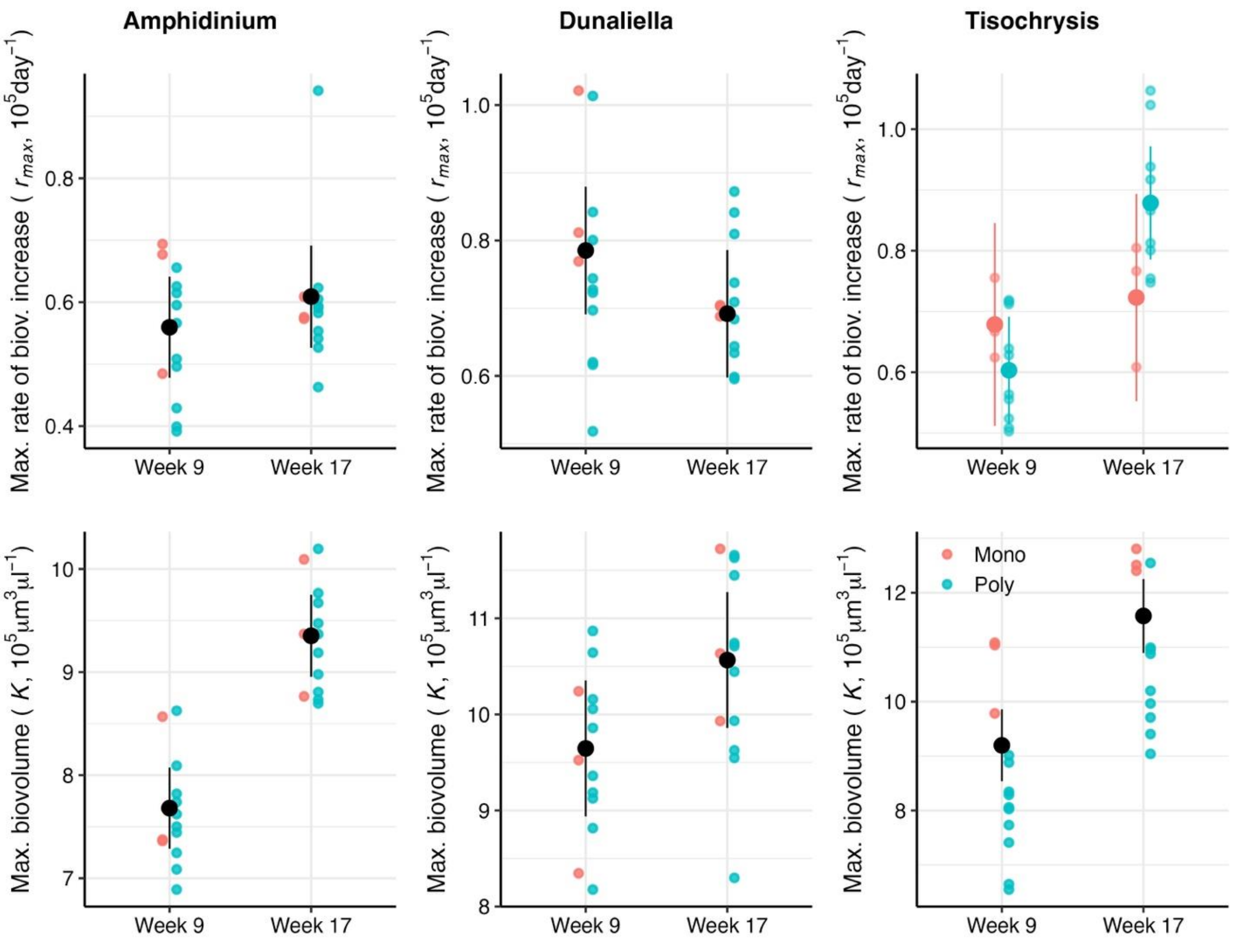

**Figure S2.** Change in maximum rate of increase ($r_{max}$) and max value (K) of biovolume ($\mu m^3$/ul) for each species after 9 and 17 weeks of evolution alone (monoculture) or in a community (polyculture). Refer to Table S2 for the model outputs.

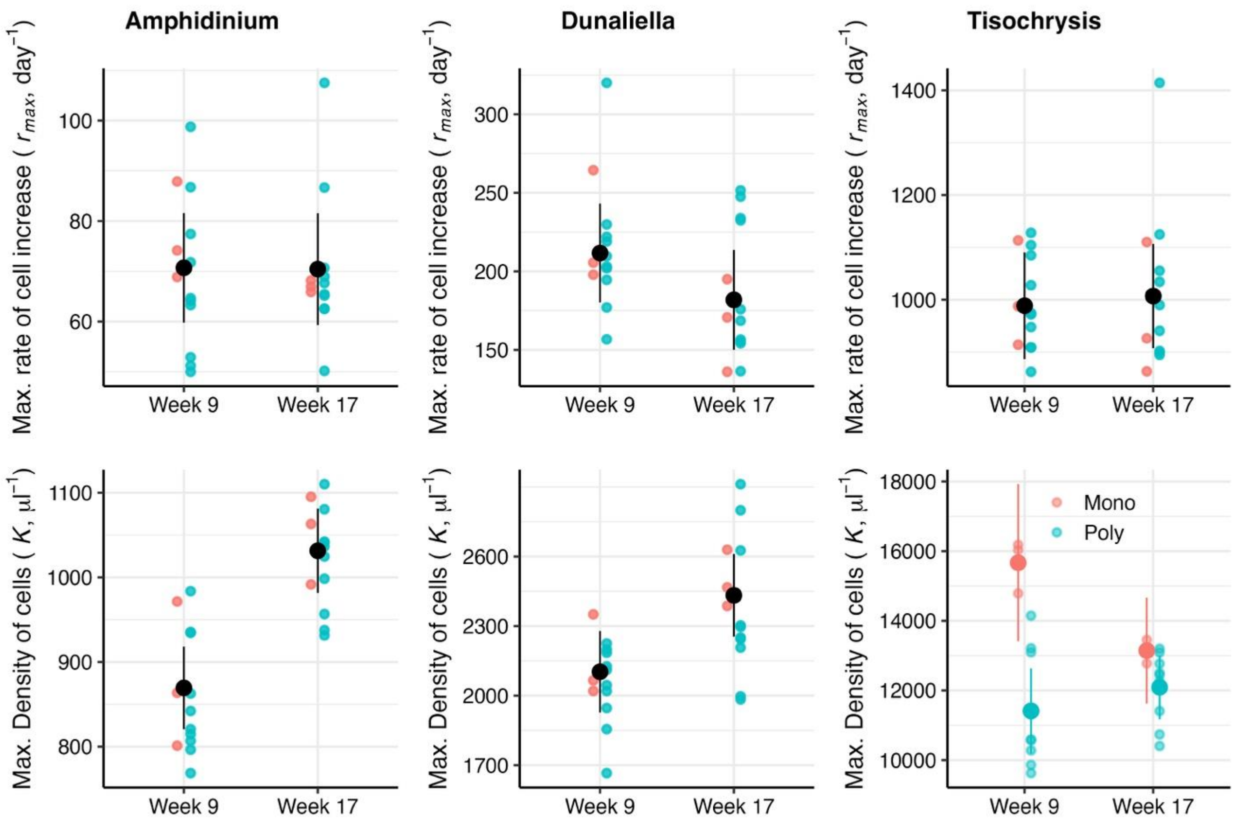

**Figure S3.** Change in maximum rate of increase ($r_{max}$) and max population density(K) (cells/μl) for each species after 9 and 17 weeks of evolution alone (monoculture) or in a community (polyculture). Refer to Table S3 for the model outputs.

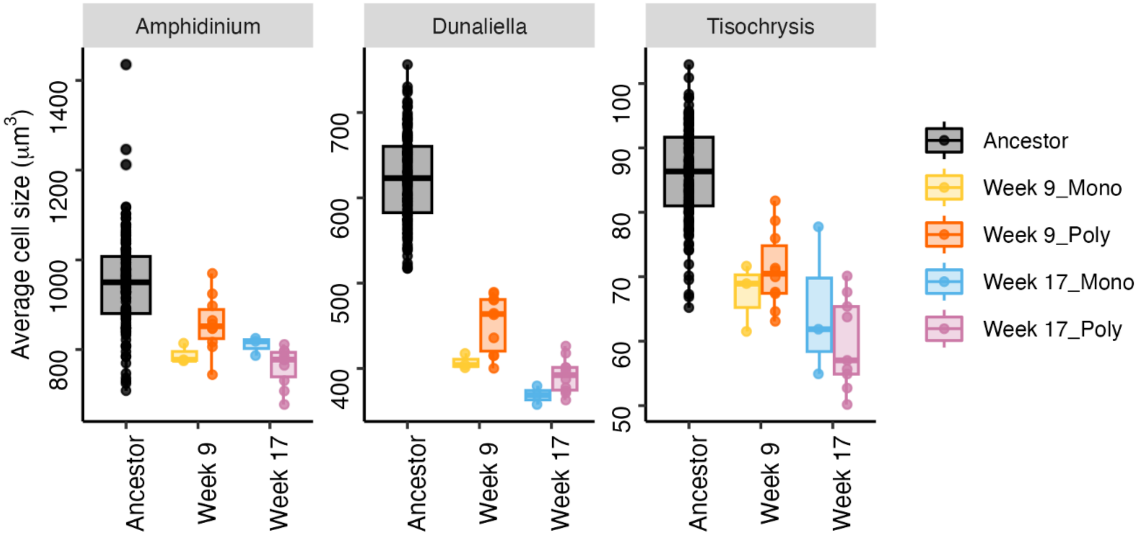

**Figure S4.** When considering only the average cell size at the start of common garden experiments (i.e. day 0, after 4 days of neutral selection), all species reduced their cell size in comparison to the ancestors. We find weak differences in size between the two common gardens (week 9 and week 17) and competition treatments. Refer to Table S4 for the model outputs.

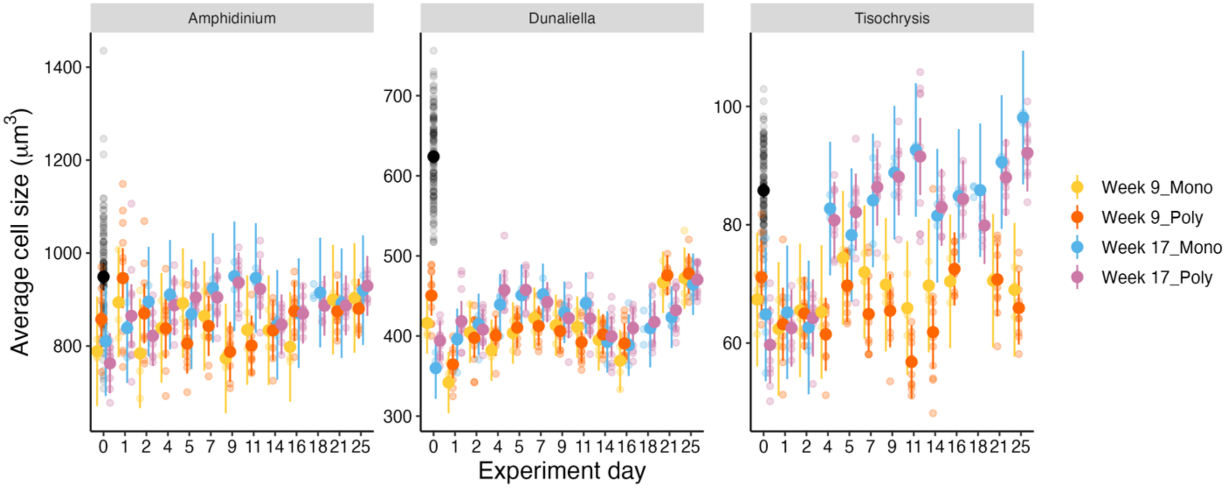

**Figure S5.** While all species reduce their cell size relative to the ancestors (black), these changes are not maintained during the common garden experiments as cell size tends to increase. This increase is particularly strong for *Tisochrysis* in the second common garden (week 17) and explains why this species increases max. biovolume without necessarily increasing cell density. Refer to Table S5 for the model outputs.

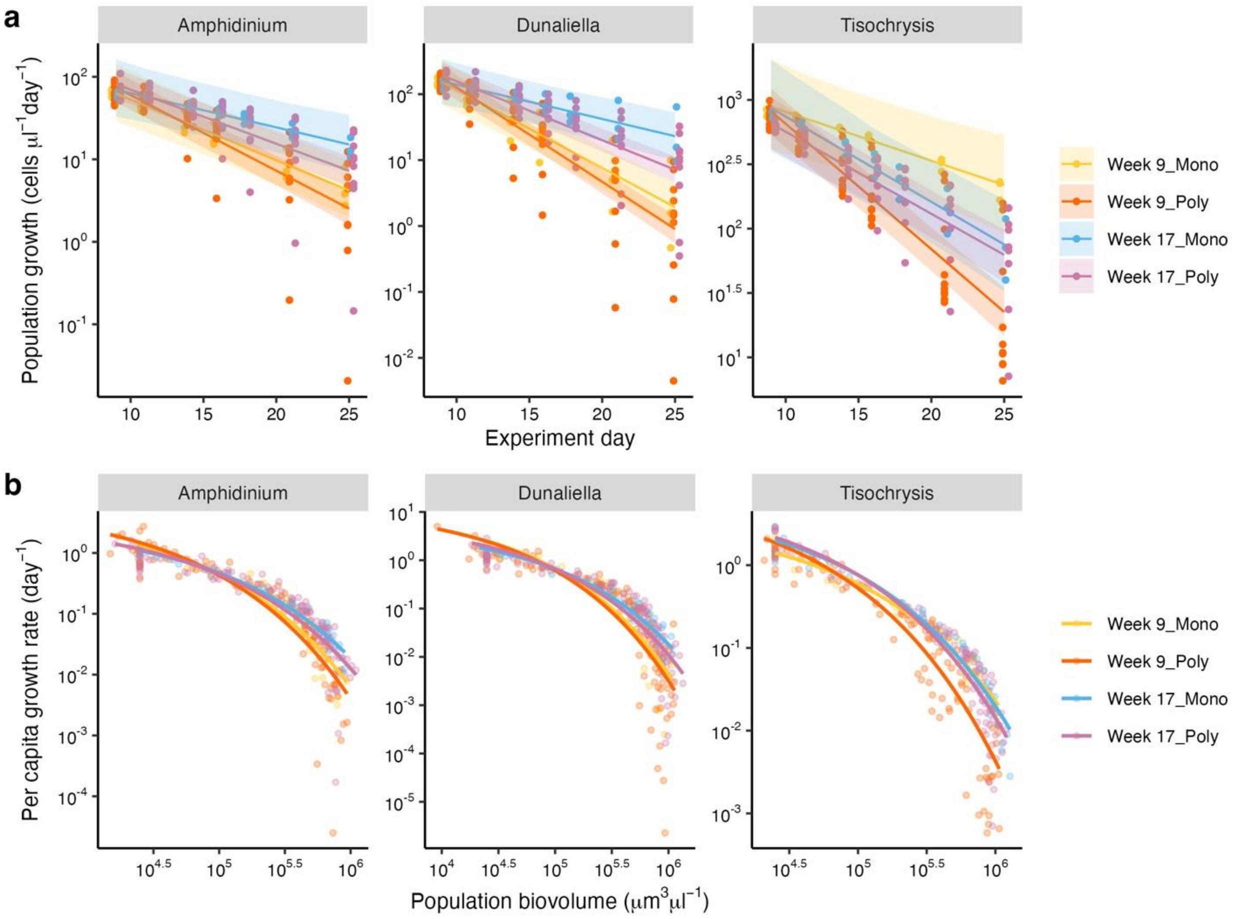

**Figure S6.** a) Population growth rate (cells μl$^{-1}$ day$^{-1}$) declines as species approach stationary phase (x-axis shows experiment day of common gardens). But this decline is shallower in week 17 compared to week 9 within each competition treatment – with one exception: *Tisochrysis* in the monoculture treatment shows the opposite pattern. Populations evolved in polyculture have stronger density-dependence than those evolved in monoculture. Refer to Table S9 for the model outputs. b) *Per capita* growth rates show the same pattern.

**Table S1**: Linear models and post-hoc test results showing the effect of competition treatment and species on the percentage change in max. growth rates ($r_{max}$) and max. values (K) of biovolume ($\mu m^3/\mu l$) and cell density (cells/µl). Relates to Figure 2. P values < 0.05 are in bold. CL = 95% confidence level. Species abbreviations are: *Dunaliella* (Dun), *Amphidinium* (Amphi) and *Tisochrysis* (Tiso). Competition treatments are mono = monoculture, poly = polyculture.

| Percentage change in the max rates of increase of biovolume (Figure 1a) | | | | | |
|---|---|---|---|---|---|
| | Df | Sum Sq | Mean Sq | F value | Pr(>F) |
| Species | 2 | 15157 | 7578.5 | 5.5275 | **0.008346** |
| Treatment | 1 | 2975 | 2975.4 | 2.1702 | 0.149909 |
| Residuals | 34 | 46616 | 1371.0 | | |
| Estimated Marginal Means | Estimate | SE | Lower CL | Upper CL | |
| Dun | -9.08 | 11.0 | -36.6 | 18.4 | |
| Amphi | 5.82 | 11.0 | -21.7 | 33.3 | |
| Tiso | 39.64 | 11.3 | 11.4 | 67.9 | |
| Contrast | Estimate | SE | Df | t ratio | p value |
| Amphi - Dun | 14.9 | 14.5 | 34 | 1.026 | 0.5659 |
| Amphi - Tiso | -33.8 | 14.8 | 34 | -2.281 | 0.0722 |
| Dun - Tiso | -48.7 | 14.8 | 34 | -3.286 | **0.0065** |

| Percentage change in max. values of biovolume (Figure 1b) | | | | | |
|---|---|---|---|---|---|
| | Df | Sum Sq | Mean Sq | F value | Pr(>F) |
| Species | 2 | 2656.3 | 1328.15 | 5.074 | **0.01221** |
| Treatment | 1 | 110.5 | 110.54 | 0.4221 | 0.52053 |
| Species × treatment | 2 | 1130.2 | 565.12 | 2.1578 | 0.13210 |
| Residuals | 32 | 8380.5 | 261.89 | | |
| Estimated Marginal Means | Estimate | SE | Lower CL | Upper CL | |
| Dun | 12.7 | 5.33 | -0.72 | 26.1 | |
| Amphi | 23.3 | 5.33 | 9.84 | 36.7 | |
| Tiso | 25.4 | 5.33 | 11.81 | 39.0 | |
| Contrast | Estimate | SE | Df | t ratio | p value |
| Amphi - Dun | 10.56 | 7.53 | 32 | 1.402 | 0.3517 |
| Amphi - Tiso | -2.14 | 7.58 | 32 | -0.282 | 0.9571 |
| Dun - Tiso | -12.70 | 7.58 | 32 | -1.676 | 0.2299 |

| Percentage change in the max rates of increase of cell density (Figure 1c) | | | | | |
|---|---|---|---|---|---|
| | Df | Sum Sq | Mean Sq | F value | Pr(>F) |
| Species | 2 | 2657 | 1328.7 | 1.2882 | 0.2889 |
| Treatment | 1 | 1703 | 1702.9 | 1.6510 | 0.2075 |
| Residuals | 34 | 35069 | 1031.4 | | |
| Estimated Marginal Means | Estimate | SE | Lower CL | Upper CL | |
| Dun | -11.29 | 9.50 | -35.1 | 12.6 | |
| Amphi | -2.91 | 9.50 | -26.8 | 20.9 | |
| Tiso | 9.58 | 9.76 | -14.9 | 34.1 | |
| Contrast | Estimate | SE | Df | t ratio | p value |
| Amphi - Dun | 8.38 | 12.6 | 34 | 0.665 | 0.7851 |
| Amphi - Tiso | -12.49 | 12.9 | 34 | -0.971 | 0.5998 |
| Dun - Tiso | -20.87 | 12.9 | 34 | -1.623 | 0.2501 |

| Percentage change in the max values of cell density (Figure 1d) | | | | | |
|---|---|---|---|---|---|
| | Df | Sum Sq | Mean Sq | F value | Pr(>F) |
| Species | 2 | 1406.2 | 703.12 | 3.6469 | **0.03743** |
| Treatment | 1 | 300.6 | 300.65 | 1.5594 | 0.22082 |
| Species × treatment | 2 | 1296.0 | 647.99 | 3.3609 | **0.04733** |
| Residuals | 32 | 6169.7 | 192.80 | | |

| Estimated Marginal Means | | | | |
|---|---|---|---|---|
| Treatment = mono | | | | |
| Species | Estimate | SE | Lower CL | Upper CL |
| Tiso | -14.8 | 8.02 | -35.027 | 5.36 |
| Dun | 18.1 | 8.02 | -2.118 | 38.27 |
| Amphi | 20.8 | 8.02 | 0.654 | 41.05 |
| Treatment = poly | | | | |
| Species | Estimate | SE | Lower CL | Upper CL |
| Tiso | 11.4 | 4.63 | -0.260 | 23.06 |
| Dun | 13.8 | 4.39 | 2.723 | 24.85 |
| Amphi | 19.3 | 4.39 | 8.192 | 30.32 |

| Contrast - Mono | Estimate | SE | Df | t ratio | p value |
|---|---|---|---|---|---|
| Amphi - Dun | 2.77 | 11.34 | 32 | 0.244 | 0.9676 |
| Amphi - Tiso | 35.68 | 11.34 | 32 | 3.147 | **0.0097** |
| Dun - Tiso | 32.91 | 11.34 | 32 | 2.903 | **0.0178** |

| Contrast - Poly | Estimate | SE | Df | t ratio | p value |
|---|---|---|---|---|---|
| Amphi - Dun | 5.47 | 6.21 | 32 | 0.881 | 0.6561 |
| Amphi - Tiso | 7.85 | 6.38 | 32 | 1.231 | 0.444 |
| Dun - Tiso | 2.38 | 6.38 | 32 | 0.374 | 0.9261 |

**Table S2**: Linear models and post-hoc test results showing the change in $r_{max}$ and maximum value (K) of biovolume ($\mu m^3/\mu l$) for each species after 9 and 17 weeks of evolution alone (mono) or in a community (poly). Initial biovolume (Biov_init) was used as a covariate. CL = 95% confidence level. Relates to Figure S2. P values < 0.05 are in bold.

| | | | | | |
|---|---|---|---|---|---|
| Species = *Amphidinium* ($r_{max}$) | | | | | |
| | Sum Sq | df | F value | Pr(>F) | |
| Intercept | 1.2140e+10 | 1 | 99.7283 | **<0.0001** | |
| Biov_init | 1.1661e+09 | 1 | 9.3679 | **0.005727** | |
| treatment | 5.9700e+07 | 1 | 0.4796 | 0.495847 | |
| Experiment | 1.5702e+08 | 1 | 1.2614 | 0.273488 | |
| Residuals | 2.7385e+09 | 22 | | | |
| Experiment | Estimate | SE | Lower CL | Upper CL | |
| Week 9 | 55959 | 3400 | 47800 | 64118 | |
| Week 17 | 60890 | 3437 | 52643 | 69137 | |
| Contrast | Estimate | SE | Df | t ratio | p value |
| Week9 – Week17 | -4931 | 4390 | 22 | -1.123 | 0.2735 |
| Species = *Amphidinium* (K) | | | | | |
| | Sum Sq | df | F value | Pr(>F) | |
| Intercept | 1.3116e+12 | 1 | 452.4749 | **<0.0001** | |
| Biov_init | 1.0241e+10 | 1 | 3.5329 | 0.07347 | |
| treatment | 8.7004e+08 | 1 | 0.3001 | 0.58931 | |
| Experiment | 1.8066e+11 | 1 | 62.3240 | **<0.0001** | |
| Residuals | 6.3772e+10 | 21 | | | |
| Experiment | Estimate | SE | Lower CL | Upper CL | |
| Week 9 | 767990 | 16408 | 728618 | 807363 | |
| Week 17 | 935249 | 16585 | 895453 | 975045 | |
| Contrast | Estimate | SE | Df | t ratio | p value |
| Week9 – Week17 | -167259 | 21187 | 22 | -7.895 | **<0.0001** |
| Species = *Dunaliella* ($r_{max}$) | | | | | |
| | Sum Sq | df | F value | Pr(>F) | |
| Intercept | 2.3852e+10 | 1 | 169.1782 | **<0.0001** | |
| Biov_init | 1.6560e+09 | 1 | 11.7454 | **0.002531** | |
| treatment | 4.3359e+08 | 1 | 3.0754 | 0.094071 | |
| Experiment | 4.2687e+08 | 1 | 3.0277 | 0.096488 | |
| Treatment × Experiment | 2.6366e+08 | 1 | 1.8701 | 0.185921 | |
| Residuals | 2.9607e+09 | 21 | | | |
| Experiment | Estimate | SE | Lower CL | Upper CL | |
| Week 17 | 69192 | 3912 | 59772 | 78613 | |
| Week 9 | 78528 | 3912 | 69109 | 87946 | |
| Contrast | Estimate | SE | Df | t ratio | p value |
| Week9 – Week17 | 9335 | 5538 | 21 | 1.686 | 0.1067 |
| Species = *Dunaliella* (K) | | | | | |
| | Sum Sq | df | F value | Pr(>F) | |
| Intercept | 3.3284e+12 | 1 | 354.8769 | **<0.0001** | |
| Biov_init | 3.8625e+10 | 1 | 4.1182 | 0.05469 | |
| treatment | 1.2582e+08 | 1 | 0.0134 | 0.90885 | |
| Experiment | 5.4894e+10 | 1 | 5.8528 | **0.02427** | |

| | | | | | |
|---|---|---|---|---|---|
| Residuals | 2.0634e+11 | 21 | | | |
| Experiment | Estimate | SE | Lower CL | Upper CL | |
| Week 9 | 94588 | 29486 | 893834 | 1035342 | |
| Week 17 | 1056584 | 29490 | 985821 | 1127347 | |
| Contrast | Estimate | SE | Df | t ratio | p value |
| Week9 – Week17 | -91996 | 38027 | 22 | -2.419 | **0.0243** |
| Species = *Tisochrysis* ($r_{max}$) | | | | | |
| | Sum Sq | df | F value | Pr(>F) | |
| Intercept | 1.1828e+10 | 1 | 113.8353 | **<0.0001** | |
| Biov_init | 1.8355e+09 | 1 | 17.6651 | **0.0004377** | |
| treatment | 1.2408e+08 | 1 | 1.1941 | 0.2874845 | |
| Experiment | 2.4711e+07 | 1 | 0.2378 | 0.6310892 | |
| Treatment × Experiment | 5.2573e+08 | 1 | 5.0597 | **0.0359108** | |
| Residuals | 2.0781e+09 | 20 | | | |
| Experiment × treatment | Estimate | SE | Lower CL | Upper CL | |
| Week 9 Poly | 60320 | 3224 | 51501 | 69138 | |
| Week 9 Mono | 67855 | 6102 | 51160 | 84549 | |
| Week 17 Mono | 72292 | 6237 | 55231 | 89353 | |
| Week 17 Poly | 87853 | 3400 | 78552 | 97155 | |
| Contrast | Estimate | SE | Df | t ratio | p value |
| Week 9 Mono – Week 17 Mono | -4437 | 9099 | 20 | -0.488 | 0.9610 |
| Week 9 Mono – Week 9 Poly | 7535 | 6895 | 20 | 1.093 | 0.6979 |
| Week 9 Mono – Week 17 Poly | -19999 | 6958 | 20 | -2.874 | **0.0428** |
| Week 17 Mono – Week 9 Poly | 119782 | 7028 | 20 | 1.703 | 0.3480 |
| Week 17 Mono – Week 17 Poly | -15562 | 7138 | 20 | -2.180 | 0.1630 |
| Week 9 Poly – Week 17 Poly | -27534 | 4684 | 20 | -5.878 | **0.0001** |
| Species = *Tisochrysis* (K) | | | | | |
| | NumDF | F value | p value | | |
| Intercept | 3.3123e+12 | 1 | 413.1748 | **<0.0001** | |
| Biov_init | 1.4355e+10 | 1 | 1.7906 | 0.1952 | |
| treatment | 2.7391e+11 | 1 | 34.1671 | **<0.0001** | |
| Experiment | 3.4033e+11 | 1 | 42.4525 | **<0.0001** | |
| Residuals | 1.6835e+11 | 21 | | | |
| Posthoc test for treatment effect | | | | | |
| | Estimate | SE | Lower CL | Upper CL | |
| Poly | 915811 | 20567 | 866289 | 965333 | |
| Mono | 1161311 | 36599 | 1073185 | 1249437 | |
| Contrast | Estimate | SE | Df | t ratio | p value |
| Mono - Poly | 245500 | 42000 | 21 | 5.845 | **<0.0001** |
| Posthoc test for Experiment effect | | | | | |
| | Estimate | SE | Lower CL | Upper CL | |
| Week9 | 919773 | 27404 | 853789 | 985757 | |
| Week17 | 1157348 | 28183 | 1089486 | 1225210 | |
| Contrast | Estimate | SE | Df | t ratio | p value |
| Week9 – Week17 | -237575 | 36463 | 21 | -6.516 | **<0.0001** |

| **Table S3**: Linear models and post-hoc test results showing the change in $r_{max}$ and maximum value (K) of cell density (cells/µl) for each species after 9 and 17 weeks of evolution alone (mono) or in a community (poly). We included initial cell density as a covariate (Density_init). For the analysis of max. cell density (K) for *Tisochrysis* we used a generalised least square model with Experiment-specific variance. CL = 95% confidence level. Relates to Figure S3. P values < 0.05 are in bold. | | | | | |
|---|---|---|---|---|---|
| Species = *Amphidinium* ($r_{max}$) | | | | | |
| | Sum Sq | df | F value | Pr(>F) | |
| Intercept | 19774.3 | 1 | 88.8262 | **<0.0001** | |
| Density_init | 1629.8 | 1 | 7.3211 | **0.01291** | |
| treatment | 28.7 | 1 | 0.1289 | 0.72297 | |
| Experiment | 0.4 | 1 | 0.0019 | 0.96592 | |
| Residuals | 4897.6 | 22 | | | |
| Experiment | Estimate | SE | Lower CL | Upper CL | |
| Week 17 | 70.4 | 4.64 | 59.3 | 81.6 | |
| Week 9 | 70.7 | 4.54 | 59.8 | 81.6 | |
| Contrast | Estimate | SE | Df | t ratio | p value |
| Week9 – Week17 | 0.256 | 5.92 | 22 | 0.043 | 0.9659 |
| Species = *Amphidinium* (K) | | | | | |
| | Sum Sq | df | F value | Pr(>F) | |
| Intercept | 2036989 | 1 | 456.9984 | **<0.0001** | |
| Density_init | 436 | 1 | 0.0978 | 0.7574 | |
| treatment | 3424 | 1 | 0.7682 | 0.3902 | |
| Experiment | 166482 | 1 | 37.3502 | **<0.0001** | |
| Resdiuals | 98061 | 22 | | | |
| Experiment | Estimate | SE | Lower CL | Upper CL | |
| Week 9 | 869 | 20.3 | 821 | 918 | |
| Week 17 | 1031 | 20.7 | 982 | 1081 | |
| Contrast | Estimate | SE | Df | t ratio | p value |
| Week9 – Week17 | -162 | 26.5 | 22 | -6.111 | **<0.0001** |
| Species = *Dunaliella* ($r_{max}$) | | | | | |
| | Sum Sq | df | F value | Pr(>F) | |
| Intercept | 204554 | 1 | 111.1146 | **<0.0001** | |
| Density_init | 7910 | 1 | 4.2968 | 0.0501 | |
| treatment | 249 | 1 | 0.1353 | 0.71650 | |
| Experiment | 5569 | 1 | 3.0250 | 0.09596 | |
| Resdiuals | 40500 | 22 | | | |
| Experiment | Estimate | SE | Lower CL | Upper CL | |
| Week 17 | 182 | 13.2 | 150 | 214 | |
| Week 9 | 212 | 13.1 | 180 | 243 | |
| Contrast | Estimate | SE | Df | t ratio | p value |
| Week9 – Week17 | 29.8 | 17.1 | 22 | 1.739 | 0.0960 |
| Species = *Dunaliella* (K) | | | | | |
| | Sum Sq | df | F value | Pr(>F) | |
| Intercept | 17091821 | 1 | 269.2690 | **<0.0001** | |
| Density_init | 135247 | 1 | 2.3444 | 0.13999 | |
| treatment | 65468 | 1 | 1.1348 | 0.29830 | |

| | | | | | |
|---|---|---|---|---|---|
| Experiment | 681788 | 1 | 11.8181 | **0.00235** | |
| Resdiuals | 1269185 | 22 | | | |
| Experiment | Estimate | SE | Lower CL | Upper CL | |
| Week 9 | 2103 | 73.3 | 1927 | 2279 | |
| Week 17 | 2433 | 74.1 | 2255 | 2611 | |
| Contrast | Estimate | SE | Df | t ratio | p value |
| Week9 – Week17 | -330 | 95.9 | 22 | -3.438 | **0.0023** |
| Species = *Tisochrysis* ($r_{max}$) | | | | | |
| | Sum Sq | df | F value | Pr(>F) | |
| Intercept | 3248627 | 1 | 193.4426 | **<0.0001** | |
| Density_init | 319240 | 1 | 19.0094 | **<0.0001** | |
| treatment | 2476 | 1 | 0.1474 | 0.7048511 | |
| Experiment | 1740 | 1 | 0.1036 | 0.7506994 | |
| Resdiuals | 352669 | 21 | | | |
| Experiment | Estimate | SE | Lower CL | Upper CL | |
| Week 9 | 988 | 42.3 | 887 | 1090 | |
| Week 17 | 1007 | 41.4 | 907 | 1107 | |
| Contrast | Estimate | SE | Df | t ratio | p value |
| Week9 – Week17 | -18.4 | 57.3 | 21 | -0.322 | 0.7507 |
| Species = *Tisochrysis* (K) | | | | | |
| | NumDF | F value | P value | | |
| Intercept | 1 | 4383.954 | **<0.0001** | | |
| Density_init | 1 | 6.581 | **0.0185** | | |
| Experiment | 1 | 1.551 | 0.2274 | | |
| Treatment | 1 | 39.257 | **<0.0001** | | |
| Experiment × treatment | 1 | 15.131 | **0.0009** | | |
| Experiment = Week 9 | Estimate | SE | Lower CL | Upper CL | |
| Mono | 11408 | 428 | 10353 | 12462 | |
| Poly | 15649 | 434 | 14130 | 17168 | |
| Experiment = Week 17 | Estimate | SE | Lower CL | Upper CL | |
| Mono | 12089 | 462 | 10966 | 13229 | |
| Poly | 13148 | 378 | 11611 | 14685 | |
| | | | cbri | | |
| Contrast | Estimate | SE | Df | t ratio | p value |
| Week 9 – Mono - Poly | 4247 | 590 | 12.6 | 7.192 | **<0.0001** |
| Week 17– Mono - Poly | 1050 | 584 | 13.7 | 1.798 | 0.0942 |

**Table S4:** Linear model and the post-hoc test results on changes in cell size between ancestors and day 0 of each common garden experiment (Week 9, Week 17). CL = 95% confidence level. P values < 0.05 are in bold. Related to Figure S4.

| Cell Size | | | | | |
|---|---|---|---|---|---|
| | Df | Sum Sq | Mean Sq | F value | Pr(>F) |
| Species | 2 | 91.113 | 45.557 | 27936.175 | **<0.0001** |
| Condition | 4 | 0.999 | 0.25 | 152.107 | **<0.0001** |
| Species × Condition | 8 | 0.137 | 0.017 | 10.422 | **<0.0001** |
| Residuals | 425 | 0.703 | 0.002 | | **<0.0001** |

Posthoc test for Experiment × species

Species = *Amphidinium*

| Experiment | Estimate | SE | Lower CL | Upper CL |
|---|---|---|---|---|
| Ancestor | 2.975 | 0.003668 | 2.968 | 2.982 |
| Week 9_Mono | 2.897 | 0.023394 | 2.851 | 2.943 |
| Week 9_Poly | 2.932 | 0.12813 | 2.907 | 2.957 |
| Week 17_Mono | 2.908 | 0.023394 | 2.862 | 2.954 |
| Week 17_Poly | 2.882 | 0.012813 | 2.857 | 2.907 |

| Contrast | Estimate | SE | Df | t ratio | p value |
|---|---|---|---|---|---|
| Ancestor – Week 9_Mono | 0.0779 | 0.0237 | 428 | 3.29 | **0.0095** |
| Ancestor – Week 9_Poly | 0.0426 | 0.0133 | 428 | 3.194 | **0.013** |
| Ancestor – Week 17_Mono | 0.0663 | 0.0237 | 428 | 2.799 | **0.0424** |
| Ancestor – Week 17_Poly | 0.093 | 0.0133 | 428 | 6.977 | **<0.0001** |
| Week 9_Mono - Week 9_Poly | -0.0353 | 0.0267 | 428 | -1.325 | 0.6758 |
| Week 9_Mono - Week 17_Mono | -0.0116 | 0.0331 | 428 | -0.351 | 0.9967 |
| Week 9_Mono - Week 17_Poly | 0.0151 | 0.0267 | 428 | 0.565 | 0.98 |
| Week 9_Poly - Week 17_Mono | 0.0237 | 0.0267 | 428 | 0.889 | 0.9009 |
| Week 9_Poly - Week 17_Poly | 0.0504 | 0.0181 | 428 | 2.782 | 0.0445 |
| Week 17_Mono - Week 17_Poly | 0.0267 | 0.0267 | 428 | 1.001 | 0.8549 |

Species = *Dunaliella*

| Experiment | Estimate | SE | Lower CL | Upper CL |
|---|---|---|---|---|
| Ancestor | 2.794 | 0.003668 | 2.787 | 2.801 |
| Week 9_Mono | 2.61 | 0.023394 | 2.564 | 2.656 |
| Week 9_Poly | 2.655 | 0.012813 | 2.63 | 2.68 |
| Week 17_Mono | 2.567 | 0.023394 | 2.521 | 2.613 |
| Week 17_Poly | 2.592 | 0.012813 | 2.567 | 2.617 |

| Contrast | Estimate | SE | Df | t ratio | p value |
|---|---|---|---|---|---|
| Ancestor – Week 9_Mono | 0.1837 | 0.0237 | 428 | 7.759 | **<0.0001** |
| Ancestor – Week 9_Poly | 0.1386 | 0.0133 | 428 | 10.396 | **<0.0001** |
| Ancestor – Week 17_Mono | 0.2272 | 0.0237 | 428 | 9.595 | **<0.0001** |
| Ancestor – Week 17_Poly | 0.2014 | 0.0133 | 428 | 15.113 | **<0.0001** |
| Week 9_Mono - Week 9_Poly | -0.0452 | 0.0267 | 428 | -1.694 | 0.439 |
| Week 9_Mono - Week 17_Mono | 0.0435 | 0.0331 | 428 | 1.314 | 0.6825 |
| Week 9_Mono - Week 17_Poly | 0.0177 | 0.0267 | 428 | 0.664 | 0.964 |
| Week 9_Poly - Week 17_Mono | 0.0887 | 0.0267 | 428 | 3.324 | **0.0085** |
| Week 9_Poly - Week 17_Poly | 0.0629 | 0.0181 | 428 | 3.47 | **0.0052** |
| Week 17_Mono - Week 17_Poly | -0.0258 | 0.0267 | 428 | -0.966 | 0.8701 |

Species = *Tisochrysis*

| Experiment | Estimate | SE | Lower CL | Upper CL |
|---|---|---|---|---|
| Ancestor | 1.932 | 0.003668 | 1.925 | 1.939 |
| Week 9_Mono | 1.828 | 0.023394 | 1.782 | 1.874 |

| | | | | |
|---|---|---|---|---|
| Week 9_Poly | 1.851 | 0.012813 | 1.826 | 1.876 |
| Week 17_Mono | 1.807 | 0.023394 | 1.761 | 1.853 |
| Week 17_Poly | 1.773 | 0.013507 | 1.747 | 1.8 |

| Contrast | Estimate | SE | Df | t ratio | p value |
|---|---|---|---|---|---|
| Ancestor – Week 9_Mono | 0.1042 | 0.0237 | 428 | 4.401 | **0.0001** |
| Ancestor – Week 9_Poly | 0.081 | 0.0133 | 428 | 6.078 | **<0.0001** |
| Ancestor – Week 17_Mono | 0.1245 | 0.0237 | 428 | 5.256 | **<0.0001** |
| Ancestor – Week 17_Poly | 0.1585 | 0.014 | 428 | 11.322 | **<0.0001** |
| Week 9_Mono - Week 9_Poly | -0.0232 | 0.0267 | 428 | -0.87 | 0.9079 |
| Week 9_Mono - Week 17_Mono | 0.0202 | 0.0331 | 428 | 0.612 | 0.9732 |
| Week 9_Mono - Week 17_Poly | 0.0543 | 0.027 | 428 | 2.009 | 0.2635 |
| Week 9_Poly - Week 17_Mono | 0.0434 | 0.0267 | 428 | 1.629 | 0.4797 |
| Week 9_Poly - Week 17_Poly | 0.0775 | 0.0186 | 428 | 4.161 | **0.0004** |
| Week 17_Mono - Week 17_Poly | 0.034 | 0.027 | 428 | 1.259 | 0.7163 |

**Table S5**: Linear models and post-hoc test results of the changes in cell size throughout each common garden experiment (week 9, week 17), competition treatment (mono, poly) and experiment day (0 to 25). Relates to Figure S5. P values < 0.05 are in bold.

| Species = *Amphidinium* | | | | | |
|---|---|---|---|---|---|
| | Df | Sum Sq | Mean Sq | F value | Pr(>F) |
| Experiment | 1 | 87824 | 87824 | 25.102 | **<0.0001** |
| Treatment | 1 | 277 | 277 | 0.0792 | 0.77862 |
| Exp_day | 12 | 202549 | 16879 | 4.8244 | **<0.0001** |
| Experiment × treatment | 1 | 6333 | 6333 | 1.8101 | 0.17961 |
| Experiment × exp_day | 11 | 326841 | 29713 | 8.4926 | **<0.0001** |
| Treatment × exp_day | 12 | 25588 | 2132 | 0.6095 | 0.83375 |
| Experiment × treatment × exp_day | 11 | 67980 | 6180 | 1.7664 | 0.05972 |
| Residuals | 274 | 958636 | 3499 | | |
| Species = *Dunaliella* | | | | | |
| | Df | Sum Sq | Mean Sq | F value | Pr(>F) |
| Experiment | 1 | 11613 | 11612.6 | 19.0158 | **<0.0001** |
| Treatment | 1 | 2699 | 2698.5 | 4.4188 | **0.03516** |
| Exp_day | 12 | 173588 | 14465.7 | 24.015 | **<0.0001** |
| Experiment × exp_day | 11 | 91494 | 8317.6 | 13.8084 | **<0.0001** |
| Treatment × exp_day | 12 | 12356 | 1029.7 | 1.7095 | 0.06418 |
| Residuals | 287 | 172877 | 602.4 | | |
| Species = *Tisochrysis* | | | | | |
| | Df | Sum Sq | Mean Sq | F value | Pr(>F) |
| Experiment | 1 | 15655.8 | 15655.8 | 488.3005 | **<0.0001** |
| Treatment | 1 | 239.9 | 239.9 | 7.481 | **0.006661** |
| Exp_day | 12 | 9172 | 764.3 | 23.8394 | **<0.0001** |
| Experiment × treatment | 1 | 30.2 | 30.2 | 0.9431 | 0.332383 |
| Experiment × exp_day | 11 | 10364.8 | 942.3 | 29.3888 | **<0.0001** |
| Treatment × exp_day | 12 | 231.6 | 19.3 | 0.6018 | 0.840007 |
| Experiment × treatment × exp_day | 11 | 463.1 | 42.1 | 1.3131 | 0.216839 |
| Residuals | 262 | 8400.2 | 32.1 | | |

**Table S6**: Linear models and post-hoc test results showing the relationship between the intrinsic rate of increase (r) and the intraspecific competition coefficient $\alpha_{ii}$ for each experiments (week 9, week 17) and competition treatments (monoculture, polyculture). r and $\alpha$ are calculated from a growth model fitted on biovolume data. CL = 95% confidence level. Relates to Figure 3a. P values < 0.05 are in bold.

| | Df | Sum Sq | Mean Sq | F value | Pr(>F) |
|---|---|---|---|---|---|
| r | 1 | 8.1790e-14 | 8.1790e-14 | 7.0151 | **0.009147** |
| Experiment | 1 | 5.7324e-13 | 5.7324e-13 | 49.1686 | **<0.0001** |
| Treatment | 1 | 1.0369e-13 | 1.0369e-13 | 8.8938 | **0.003779** |
| species | 2 | 9.9598e-13 | 4.9799e-13 | 42.7143 | **<0.0001** |
| r × species | 2 | 4.5700e-14 | 2.2850e-14 | 1.96 | 0.147487 |
| Experiment × treatment | 1 | 1.7200e-14 | 1.7200e-14 | 1.4749 | 0.228102 |
| Experiment × species | 2 | 1.0862e-13 | 5.4310e-14 | 4.6582 | **0.012164** |
| Treatment × species | 2 | 4.5510e-14 | 2.2750e-14 | 1.9517 | 0.148648 |
| Residuals | 81 | 9.4435e-13 | 1.1660e-14 | | |

Posthoc test for Experiment × species

Species = *Amphidinium*

| Experiment | Estimate | SE | Lower CL | Upper CL |
|---|---|---|---|---|
| Week 9 | 1.40e-06 | 3.17e-08 | 1.33e-06 | 1.46e-06 |
| Week 17 | 1.20e-06 | 2.79e-08 | 1.15e-06 | 1.26e-06 |

Species = *Dunaliella*

| Experiment | Estimate | SE | Lower CL | Upper CL |
|---|---|---|---|---|
| Week 9 | 1.11e-06 | 2.98e-08 | 1.05e-06 | 1.17e-06 |
| Week 17 | 1.04e-06 | 2.76e-08 | 9.87e-07 | 1.10e-06 |

Species = *Tisochrysis*

| Experiment | Estimate | SE | Lower CL | Upper CL |
|---|---|---|---|---|
| Week 9 | 1.18e-06 | 2.85e-08 | 1.12e-06 | 1.23e-06 |
| Week 17 | 9.54e-07 | 2.83e-08 | 8.97e-07 | 1.01e-06 |

Species =*Amphidinium*

| Contrast | Estimate | SE | Df | t ratio | p value |
|---|---|---|---|---|---|
| Week9 – Week17 | 1.97e-07 | 4.33e-08 | 81 | 4.555 | **<0.0001** |

Species =*Dunaliella*

| Contrast | Estimate | SE | Df | t ratio | p value |
|---|---|---|---|---|---|
| Week9 – Week17 | 6.86e-08 | 4.06e-08 | 81 | 1.691 | 0.0948 |

Species =*Tisochrysis*

| Contrast | Estimate | SE | Df | t ratio | p value |
|---|---|---|---|---|---|
| Week9 – Week17 | 2.24e-07 | 3.95e-08 | 81 | 5.664 | **<0.0001** |

Posthoc test for Competition Treatment

| Treatment | Estimate | SE | Lower CL | Upper CL |
|---|---|---|---|---|
| Mono | 1.12e-06 | 1.93e-08 | 1.08e-06 | 1.16e-06 |
| Poly | 1.18e-06 | 1.43e-08 | 1.15e-06 | 1.20e-06 |

Competition Treatment

| Contrast | Estimate | SE | Df | t ratio | p value |
|---|---|---|---|---|---|
| Mono - Poly | -5.78e-08 | 2.46e-08 | 81 | -2.349 | **0.0213** |

**Table S7**: Linear mixed effect models and post-hoc test on changes in *per capita* net energy production (J/day) with biovolume (biov; µm$^3$/µl) (both $log_{10}$-transformed) between competition treatments (mono, poly), experiments (week9, week 17) and species (*Amphidinium*, *Dunaliella*, *Tisochrysis*). CL = 95% confidence level. P values < 0.05 are in bold. Related to Figure 3b.

| | Sum Sq | Mean Sq | DF | DenDF | F value | Pr(>F) |
|---|---|---|---|---|---|---|
| $log_{10}$(biov) | 10.6951 | 10.6951 | 1 | 644.57 | 414.1516 | **<0.0001** |
| Experiment | 2.5178 | 2.5178 | 1 | 646.64 | 97.4962 | **<0.0001** |
| treatment | 0.01287 | 0.0128 | 1 | 646.64 | 0.4964 | 0.481334 |
| species | 0.0909 | 0.0455 | 2 | 646.57 | 1.7606 | 0.172772 |
| $log_{10}$ (biov) × Experiment | 2.7850 | 2.7850 | 1 | 644.57 | 107.8456 | **<0.0001** |
| $log_{10}$ (biov) × treatment | 0.0153 | 0.0153 | 1 | 644.57 | 0.5917 | 0.442051 |
| Experiment × treatment | 0.0866 | 0.0866 | 1 | 646.64 | 3.3535 | 0.067523 |
| $log_{10}$ (biov) × species | 0.3232 | 0.1616 | 2 | 644.49 | 6.2581 | **0.002033** |
| Experiment × species | 0.5247 | 0.2624 | 2 | 646.57 | 10.1598 | **<0.0001** |
| Treatment × species | 0.0305 | 0.0153 | 2 | 646.57 | 0.5910 | 0.554091 |
| $log_{10}$ (biov) × Experiment × treatment | 0.0750 | 0.0750 | 1 | 644.50 | 2.9030 | *0.088896* |
| $log_{10}$ (biov) × Experiment × species | 0.5764 | 0.2882 | 2 | 644.50 | 11.1606 | **<0.001** |
| $log_{10}$ (biov) × treatment × species | 0.0290 | 0.0145 | 2 | 644.49 | 0.5621 | 0.570292 |
| Experiment × treatment × species | 0.0825 | 0.0413 | 2 | 646.57 | 1.5976 | 0.203174 |
| $log_{10}$(biov) × Experiment × treatment × species | 0.0803 | 0.0402 | 2 | 644.50 | 1.5553 | 0.211911 |

Posthoc test for biovolume × Experiment × Species

| Species | Experiment | Slope estimates and CL |
|---|---|---|
| *Amphidinium* | Week 9 | -0.979 (-1.124, -0.8344) |
| | Week 17 | -0.0508 (-0.192, -0.0907) |
| *Dunaliella* | Week 9 | -0.7601 (-0.918, -0.6027) |
| | Week 17 | -0.5205 (-0.677, -0.3641) |
| *Tisochrysis* | Week 9 | -1.1943 (-1.350, -1.0389) |
| | Week 17 | -0.3801 (-0.542, -0.2183) |

Species = *Amphidinium*

| Contrast | Estimate | SE | Df | t ratio | p value |
|---|---|---|---|---|---|
| Week9 – Week17 | -0.929 | 0.103 | 644 | -9.000 | **<0.0001** |

Species = *Dunaliella*

| Contrast | Estimate | SE | Df | t ratio | p value |
|---|---|---|---|---|---|
| Week9 – Week17 | -0.270 | 0.113 | 644 | -2.121 | **0.0343** |

Species = *Tisochrysis*

| Contrast | Estimate | SE | Df | t ratio | p value |
|---|---|---|---|---|---|
| Week9 – Week17 | -0.814 | 0.114 | 650 | -7.127 | **<0.0001** |

Posthoc test for biovolume × Experiment × Treatment (p = 0.09)

| Experiment | Competition treatment | Slope estimates and CL |
|---|---|---|
| Week 9 | Mono | -1.008 (-1.166, -0.8497) |
| | Poly | -0.948 (-1.027, -0.8699) |

| | | | | | |
|---|---|---|---|---|---|
| Week 17 | | Mono<br>Poly | | -0.238 (-0.393, -0.0835)<br>-0.396 (-0.482, -0.3099) | |
| Experiment = Week 9 | | | | | |
| Contrast | Estimate | SE | Df | t ratio | p value |
| Mono – Poly | -0.0595 | 0.0899 | 641 | -0.662 | 0.5083 |
| Experiment = Week 17 | | | | | |
| Contrast | Estimate | SE | Df | t ratio | p value |
| Mono – Poly | 0.1574 | 0.0902 | 651 | 1.744 | 0.0816 |

**Table S8**: Linear model and the post-hoc test results on changes in *per capita* photosynthesis and respiration rates (J/min, $log_{10}$-transformed) between species, condition (experiment and competition treatment combined) and phase (day 4 in the middle of exponential phase vs day 25 in stationary phase). Related to Figure 4. CL = 95% confidence level. P values < 0.05 are in bold.

| Photosynthesis | | | | | |
|---|---|---|---|---|---|
| | Df | Sum Sq | Mean Sq | F value | Pr(>F) |
| Species | 2 | 21.7368 | 10.8684 | 441.456 | **<0.001** |
| Condition (experiment and treatment combined) | 3 | 0.8226 | 0.2742 | 11.137 | **<0.001** |
| Phase (Day 4 vs Day 25) | 1 | 4.8380 | 4.8380 | 196.5127 | **<0.001** |
| Species × condition | 6 | 0.0688 | 0.0115 | 0.4658 | 0.832596 |
| Species × phase | 2 | 0.2454 | 0.1227 | 4.49838 | **0.008214** |
| Condition × phase | 3 | 0.3178 | 0.1059 | 4.3023 | **0.006268** |
| Species × condition × phase | 6 | 0.3126 | 0.0521 | 2.1161 | **0.055633** |
| Residuals | 130 | 3.2005 | 0.0246 | | |

| Posthoc test for species × condition × phase | | | | |
|---|---|---|---|---|
| Species = *Amphidinium*, Phase = Exponential | | | | |
| Condition | Estimate | SE | Lower CL | Upper CL |
| Week 9_Mono | -8.31 | 0.0906 | -8.48 | -8.13 |
| Week 9_Poly | -8.25 | 0.0496 | -8.35 | -8.15 |
| Week 17_Mono | -8.29 | 0.0906 | -8.46 | -8.11 |
| Week 17_Poly | -8.15 | 0.0496 | -8.25 | -8.06 |
| Species = *Amphidinium*, Phase = Stationary | | | | |
| Condition | Estimate | SE | Lower CL | Upper CL |
| Week 9_Mono | -8.59 | 0.0906 | -8.77 | -8.41 |
| Week 9_Poly | -8.63 | 0.0496 | -8.73 | -8.54 |
| Week 17_Mono | -8.41 | 0.0906 | -8.59 | -8.23 |
| Week 17_Poly | -8.4 | 0.0496 | -8.49 | -8.3 |
| Species = *Dunaliella*, Phase = Exponential | | | | |
| Condition | Estimate | SE | Lower CL | Upper CL |
| Week 9_Mono | -8.59 | 0.0906 | -8.77 | -8.41 |
| Week 9_Poly | -8.59 | 0.0496 | -8.69 | -8.49 |
| Week 17_Mono | -8.5 | 0.0906 | -8.68 | -8.32 |
| Week 17_Poly | -8.4 | 0.0496 | -8.5 | -8.3 |
| Species = *Dunaliella*, Phase = Stationary | | | | |
| Condition | Estimate | SE | Lower CL | Upper CL |
| Week 9_Mono | -8.9 | 0.0906 | -9.08 | -8.72 |
| Week 9_Poly | -8.92 | 0.0496 | -9.02 | -8.82 |
| Week 17_Mono | -8.76 | 0.0906 | -8.94 | -8.58 |
| Week 17_Poly | -8.71 | 0.0496 | -8.81 | -8.61 |
| Species = *Tisochrysis*, Phase = Exponential | | | | |
| Condition | Estimate | SE | Lower CL | Upper CL |
| Week 9_Mono | -8.9 | 0.0906 | -9.08 | -8.72 |
| Week 9_Poly | -9.03 | 0.0496 | -9.13 | -8.93 |
| Week 17_Mono | -9.11 | 0.0906 | -9.29 | -8.93 |
| Week 17_Poly | -9.08 | 0.0523 | -9.18 | -8.97 |
| Species = *Tisochrysis*, Phase = Stationary | | | | |
| Condition | Estimate | SE | Lower CL | Upper CL |
| Week 9_Mono | -9.81 | 0.0906 | -9.99 | -9.63 |
| Week 9_Poly | -9.58 | 0.0496 | -9.68 | -9.49 |
| Week 17_Mono | -9.4 | 0.0906 | -9.58 | -9.22 |

| | | | | | |
|---|---|---|---|---|---|
| Week 17_Poly | -9.36 | 0.0523 | -9.47 | -9.26 | |
| Phase = exponential, species = *Amphidinium* | | | | | |
| Contrast | Estimate | SE | Df | t ratio | p value |
| Week 9_Mono - Week 9_Poly | -0.05312 | 0.1033 | 130 | -0.514 | 0.9556 |
| Week 9_Mono - Week 17_Mono | -0.0206 | 0.1281 | 130 | -0.161 | 0.9985 |
| Week 9_Mono - Week 17_Poly | -0.15113 | 0.1033 | 130 | -1.463 | 0.4627 |
| Week 9_Poly - Week 17_Mono | 0.03252 | 0.1033 | 130 | 0.315 | 0.9891 |
| Week 9_Poly - Week 17_Poly | -0.09801 | 0.0702 | 130 | -1.397 | 0.5037 |
| Week 17_Mono - Week 17_Poly | -0.13053 | 0.1033 | 130 | -1.264 | 0.5876 |
| Phase = stationary, species = *Amphidinium* | | | | | |
| Contrast | Estimate | SE | Df | t ratio | p value |
| Week 9_Mono - Week 9_Poly | 0.04381 | 0.1033 | 130 | 0.424 | 0.9743 |
| Week 9_Mono - Week 17_Mono | -0.1826 | 0.1281 | 130 | -1.425 | 0.4859 |
| Week 9_Mono - Week 17_Poly | -0.19459 | 0.1033 | 130 | -1.884 | 0.24 |
| Week 9_Poly - Week 17_Mono | -0.22642 | 0.1033 | 130 | -2.192 | 0.1309 |
| Week 9_Poly - Week 17_Poly | -0.23841 | 0.0702 | 130 | -3.398 | **0.0049** |
| Week 17_Mono - Week 17_Poly | -0.01199 | 0.1033 | 130 | -0.116 | 0.9994 |
| Phase = exponential, species = *Dunaliella* | | | | | |
| Contrast | Estimate | SE | Df | t ratio | p value |
| Week 9_Mono - Week 9_Poly | 0.00455 | 0.1033 | 130 | 0.044 | 1 |
| Week 9_Mono - Week 17_Mono | -0.0875 | 0.1281 | 130 | -0.683 | 0.9034 |
| Week 9_Mono - Week 17_Poly | -0.18613 | 0.1033 | 130 | -1.802 | 0.2771 |
| Week 9_Poly - Week 17_Mono | -0.09205 | 0.1033 | 130 | -0.891 | 0.8094 |
| Week 9_Poly - Week 17_Poly | -0.19068 | 0.0702 | 130 | -2.717 | **0.0371** |
| Week 17_Mono - Week 17_Poly | -0.09863 | 0.1033 | 130 | -0.955 | 0.7752 |
| Phase = stationary, species = *Dunaliella* | | | | | |
| Contrast | Estimate | SE | Df | t ratio | p value |
| Week 9_Mono - Week 9_Poly | 0.02427 | 0.1033 | 130 | 0.235 | 0.9954 |
| Week 9_Mono - Week 17_Mono | -0.1377 | 0.1281 | 130 | -1.075 | 0.7055 |
| Week 9_Mono - Week 17_Poly | -0.18782 | 0.1033 | 130 | -1.818 | 0.2694 |
| Week 9_Poly - Week 17_Mono | -0.16197 | 0.1033 | 130 | -1.568 | 0.4004 |
| Week 9_Poly - Week 17_Poly | -0.21208 | 0.0702 | 130 | -3.022 | **0.0158** |
| Week 17_Mono - Week 17_Poly | -0.05011 | 0.1033 | 130 | -0.485 | 0.9623 |
| Phase = exponential, species = *Tisochrysis* | | | | | |
| Contrast | Estimate | SE | Df | t ratio | p value |
| Week 9_Mono - Week 9_Poly | 0.13353 | 0.1033 | 130 | 1.293 | 0.5691 |
| Week 9_Mono - Week 17_Mono | 0.21231 | 0.1281 | 130 | 1.657 | 0.3506 |
| Week 9_Mono - Week 17_Poly | 0.18131 | 0.1046 | 130 | 1.733 | 0.3108 |
| Week 9_Poly - Week 17_Mono | 0.07878 | 0.1033 | 130 | 0.763 | 0.871 |
| Week 9_Poly - Week 17_Poly | 0.04777 | 0.0721 | 130 | 0.663 | 0.9109 |
| Week 17_Mono - Week 17_Poly | -0.03101 | 0.1046 | 130 | -0.296 | 0.9909 |
| Phase = stationary, species = *Tisochrysis* | | | | | |
| Contrast | Estimate | SE | Df | t ratio | p value |
| Week 9_Mono - Week 9_Poly | -0.22491 | 0.1033 | 130 | -2.178 | 0.135 |
| Week 9_Mono - Week 17_Mono | -0.40739 | 0.1281 | 130 | -3.18 | **0.0098** |
| Week 9_Mono - Week 17_Poly | -0.44544 | 0.1046 | 130 | -4.258 | **0.0002** |
| Week 9_Poly - Week 17_Mono | -0.18248 | 0.1033 | 130 | -1.767 | 0.2941 |
| Week 9_Poly - Week 17_Poly | -0.22053 | 0.0721 | 130 | -3.059 | **0.0142** |
| Week 17_Mono - Week 17_Poly | -0.03804 | 0.1046 | 130 | -0.364 | 0.9835 |

| Respiration | | | | | |
|---|---|---|---|---|---|
| | Df | Sum Sq | Mean Sq | F value | Pr(>F) |
| Species | 2 | 35.964 | 17.819 | 176.1180 | **<0.0001** |
| Condition (experiment and treatment combined) | 3 | 1.418 | 0.4727 | 4.6295 | **0.004117** |
| Phase (Day 4 vs Day 25) | 1 | 1.673 | 1.6727 | 16.3830 | **<0.0001** |
| Condition × phase | 3 | 1.127 | 0.3758 | 3.6802 | **0.013822** |
| Residuals | 132 | 13.477 | 0.1021 | | |
| Posthoc test for condition × phase | | | | | |
| Phase = Exponential | | | | | |
| Condition | Estimate | SE | Lower CL | Upper CL | |
| Week 9_Mono | -9.55 | 0.1065 | -9.76 | -9.34 | |
| Week 9_Poly | -9.52 | 0.0583 | -9.63 | -9.4 | |
| Week 17_Mono | -9.73 | 0.1209 | -9.97 | -9.49 | |
| Week 17_Poly | -9.98 | 0.0743 | -10.13 | -9.84 | |
| Phase = Stationary | | | | | |
| Condition | Estimate | SE | Lower CL | Upper CL | |
| Week 9_Mono | -9.54 | 0.1065 | -9.75 | -9.33 | |
| Week 9_Poly | -9.43 | 0.0583 | -9.54 | -9.31 | |
| Week 17_Mono | -9.48 | 0.1065 | -9.69 | -9.26 | |
| Week 17_Poly | -9.52 | 0.0594 | -9.64 | -9.40 | |
| Phase = Exponential | | | | | |
| Contrast | Estimate | SE | Df | t ratio | p value |
| Week 9_Mono - Week 9_Poly | -0.0344 | 0.1214 | 132 | -0.283 | 0.992 |
| Week 9_Mono - Week 17_Mono | 0.1835 | 0.1611 | 132 | 1.139 | 0.6662 |
| Week 9_Mono - Week 17_Poly | 0.4333 | 0.1299 | 132 | 3.336 | **0.006** |
| Week 9_Poly - Week 17_Mono | 0.2179 | 0.1342 | 132 | 1.623 | 0.3692 |
| Week 9_Poly - Week 17_Poly | 0.4677 | 0.0945 | 132 | 4.95 | **<0.0001** |
| Week 17_Mono - Week 17_Poly | 0.2498 | 0.1419 | 132 | 1.76 | 0.2975 |
| Phase = Stationary | | | | | |
| Contrast | Estimate | SE | Df | t ratio | p value |
| Week 9_Mono - Week 9_Poly | -0.1158 | 0.1214 | 132 | -0.953 | 0.776 |
| Week 9_Mono - Week 17_Mono | -0.0669 | 0.1506 | 132 | -0.444 | 0.9707 |
| Week 9_Mono - Week 17_Poly | -0.0212 | 0.1219 | 132 | -0.174 | 0.9981 |
| Week 9_Poly - Week 17_Mono | 0.0489 | 0.1214 | 132 | 0.403 | 0.9778 |
| Week 9_Poly - Week 17_Poly | 0.0946 | 0.0832 | 132 | 1.137 | 0.6674 |
| Week 17_Mono - Week 17_Poly | 0.0457 | 0.1219 | 132 | 0.375 | 0.982 |
| Posthoc test for main species effect | | | | | |
| Species | Estimate | SE | Lower CL | Upper CL | |
| *Amphidinium* | -9.01 | 0.0515 | -9.11 | -8.91 | |
| *Dunaliella* | -9.54 | 0.0484 | -9.63 | -9.44 | |
| *Tisochrysis* | -10.23 | 0.0492 | -10.33 | -10.14 | |
| Contrast | Estimate | SE | Df | t ratio | p value |
| *Amphidinium - Dunaliella* | 0.526 | 0.0655 | 132 | 7.905 | **<0.0001** |
| *Amphidinium - Tisochrysis* | 1.223 | 0.0673 | 132 | 18.182 | **<0.0001** |
| *Dunaliella - Tisochrysis* | 0.697 | 0.0649 | 132 | 10.740 | **<0.0001** |

**Table S9**: Linear Mixed-Effects Model and post-hoc tests for changes in the population growth (cells per day; $log_{10}$-transformed) in the declining phase (day 8 onwards) considering the effect of experiment day (8 to 25), experiment (Week 9, Week 17), competition treatment (mono, poly) and species (*Amphidinium, Dunaliella, Tisochrysis*). We included the unique sample ID as random effect. CL = 95% confidence level. Relates to Figure S6. P values < 0.05 are in bold.

Species = *Amphidinium*

| | Sum Sq | Mean Sq | NumDF | DenDF | F value | Pr(>F) |
|---|---|---|---|---|---|---|
| Exp_day | 62.887 | 62.887 | 1 | 410.47 | 1063.673 | **<0.0001** |
| Experiment | 0.393 | 0.393 | 1 | 241.28 | 6.644 | **0.010544** |
| Treatment | 0.372 | 0.372 | 1 | 241.28 | 6.2845 | **0.012838** |
| Species | 3.846 | 1.923 | 2 | 241.28 | 32.5302 | **<0.0001** |
| Exp_day × Experiment | 2.426 | 2.426 | 1 | 410.47 | 41.0396 | **<0.0001** |
| Exp_day × treatment | 1.866 | 1.866 | 1 | 410.47 | 31.5665 | **<0.0001** |
| Experiment × treatment | 0.026 | 0.026 | 1 | 241.28 | 0.4361 | 0.509634 |
| Exp_day × species | 2.649 | 1.325 | 2 | 410.47 | 22.4054 | **<0.0001** |
| Experiment × species | 0.173 | 0.087 | 2 | 241.28 | 1.4666 | 0.232763 |
| Treatment × species | 0.026 | 0.013 | 2 | 241.28 | 0.2226 | 0.800603 |
| Exp_day × Experiment × treatment | 0.127 | 0.127 | 1 | 410.47 | 2.14 | 0.144268 |
| Exp_day × Experiment × species | 1.416 | 0.708 | 2 | 410.47 | 11.9737 | **<0.0001** |
| Exp_day × treatment × species | 0.041 | 0.020 | 2 | 410.47 | 0.3453 | 0.708196 |
| Experiment × treatment × species | 0.162 | 0.081 | 2 | 241.28 | 1.3734 | 0.255216 |
| Exp_day × Experiment × treatment × species | 0.737 | 0.369 | 2 | 410.47 | 6.2346 | **0.002151** |

treatment = Mono, species = *Amphidinium*:

| | Estimate | SE | Lower CL | Upper CL | |
|---|---|---|---|---|---|
| Week 9 | -0.0730 | 0.01035 | -0.0933 | -0.0526 | |
| Week 17 | -0.0407 | 0.01025 | -0.0609 | -0.0205 | |
| contrast | estimate | SE | df | t ratio | p value |
| Week 9 – Week 17 | -0.0323 | 0.01457 | 411 | -2.215 | **0.0273** |

treatment = Poly, species = *Amphidinium*:

| | Estimate | SE | Lower CL | Upper CL | |
|---|---|---|---|---|---|
| Week 9 | -0.0913 | 0.00567 | -0.1025 | -0.0802 | |
| Week 17 | -0.0653 | 0.00562 | -0.0763 | -0.0542 | |
| contrast | estimate | SE | df | t ratio | p value |
| Week 9 – Week 17 | -0.0260 | 0.00798 | 411 | -3.263 | **0.0012** |

treatment = Mono, species = *Dunaliella*:

| | Estimate | SE | Lower CL | Upper CL | |
|---|---|---|---|---|---|
| Week 9 | -0.1193 | 0.01035 | -0.1397 | -0.0990 | |
| Week 17 | -0.0522 | 0.01025 | -0.0723 | -0.0320 | |
| contrast | estimate | SE | df | t ratio | p value |
| Week 9 – Week 17 | -0.0672 | 0.01457 | 411 | -4.610 | **<0.0001** |

treatment = Poly, species = *Dunaliella*:

| | Estimate | SE | Lower CL | Upper CL | |
|---|---|---|---|---|---|
| Week 9 | -0.1432 | 0.00567 | -0.1544 | -0.1321 | |
| Week 17 | -0.0896 | 0.00562 | -0.1006 | -0.0786 | |
| contrast | estimate | SE | df | t ratio | p value |
| Week 9 – Week 17 | -0.0536 | 0.00798 | 411 | -6.718 | **<0.0001** |

| | | | | | |
|---|---|---|---|---|---|
| treatment = Mono, species = *Tisochrysis*: | | | | | |
| | Estimate | SE | Lower CL | Upper CL | |
| Week 9 | -0.0379 | 0.01035 | -0.0583 | -0.0176 | |
| Week 17 | -0.0661 | 0.01025 | -0.0863 | -0.0459 | |
| contrast | estimate | SE | df | t ratio | p value |
| Week 9 – Week 17 | 0.0282 | 0.01457 | 411 | 1.934 | 0.0538 |
| treatment = Poly, species = *Tisochrysis*: | | | | | |
| | Estimate | SE | Lower CL | Upper CL | |
| Week 9 | -0.0978 | 0.00567 | -0.1089 | -0.0866 | |
| Week 17 | -0.0640 | 0.00592 | -0.0756 | -0.0523 | |
| contrast | estimate | SE | df | t ratio | p value |
| Week 9 – Week 17 | -0.0338 | 0.00820 | 411 | -4.123 | **<0.0001** |
